# The Role of Magnetic and Celestial Cues in Orientation and Navigation of Red Underwing (*Catocala nupta),* a European Migratory Moth

**DOI:** 10.64898/2026.03.04.709557

**Authors:** Aleksandr Pakhomov, David Dreyer, Thomas Zechmeister, Henrik Mouritsen, Dmitry Kishkinev

## Abstract

Nocturnal migration is a remarkable phenomenon observed in many insect species, including moths. Migratory moths are capable of maintaining precise directional orientation during migration, as demonstrated in both laboratory and field studies, suggesting that they use multiple environmental cues for orientation and navigation. Recent studies on Australian Bogong moths revealed that these animals can use stellar cues and likely the geomagnetic field (in conjunction with local visual cues) to select and maintain population-specific migratory direction. However, the underlying orientation mechanisms used by most other migratory moths are still largely unresolved. Further, it remains unclear whether migratory moths can adjust their orientation using Earth’s magnetic field parameters for determining their position relative to the goal (i.e. location or map information) – an ability clearly shown in some migratory birds which respond to virtual magnetic displacements by correcting their orientation (experiments when animals are exposed to magnetic cues corresponding to other geographic locations).

Here, we present results from virtual magnetic displacement experiments conducted on red underwings (*Catocala nupta*). In addition, we tested their orientation under simulated overcast conditions and in a vertical magnetic field to get indications whether this species relies on geomagnetic or celestial cues to maintain its population-specific migratory direction. Our results show that (1) red underwings did not compensate for virtual magnetic displacement, indicating the absence of a magnetic map; (2) they remained significantly oriented in the absence of geomagnetic information, suggesting the use of a stellar compass; and (3) there was no evidence of magnetic compass orientation in absence of any visual cues.

## Introduction

Nocturnal migration is a remarkable phenomenon observed in many insect species including moths. These insects can maintain a precise direction during migration, as shown in both laboratory and field studies (Chapman et al., 2008; Chapman et al., 2010; Dreyer et al., 2018a; Menz et al., 2022; Chen et al., 2023; Gao et al., 2024; Werber et al., 2025; Dreyer et al., 2025), indicating their ability to use multiple environmental cues for orientation (direction-finding, compass systems) and/or navigation (position-finding, maps). However, the exact underlying orientation mechanisms that enable them to select and maintain a seasonally and population appropriate direction towards their migratory goal during night-time flight (compass systems) remain poorly understood. While over 50 years of extensive research on night-migrating birds has demonstrated the importance of celestial- and geomagnetic-based orientation mechanisms in these animals (time-independent star compass and light-dependent inclination magnetic compass, (Wiltschko and Wiltschko, 1995; Wiltschko and Wiltschko, 2015; Mouritsen, 2018; Mouritsen, 2022), comparatively little is known about how these mechanisms function in nocturnal insects.

Among Lepidopteran migrants, detailed studies of orientation mechanisms have been conducted using only a handful of iconic species: the diurnal North American monarch butterfly (*Danaus plexippus*) (Urquhart and Urquhart, 1978; Mouritsen and Frost, 2002a; Stalleicken et al., 2005; Mouritsen et al., 2013a; Guerra and Reppert, 2015; Beetz et al., 2021; Beetz et al., 2023; Beetz and el Jundi, 2023) and the nocturnal Australian Bogong moth (*Agrotis infusa*) (Warrant et al., 2016; Dreyer et al., 2018a; Green et al., 2021; Dreyer et al., 2025). As diurnal migrants, monarch butterflies use a time-compensated sun compass and it has been suggested that they also use a light-dependent inclination magnetic compass [Guerra et al., 2014, but see Stalleicken et al. 2005], similar to birds (Ritz et al., 2000; Wiltschko and Wiltschko, 2015; Hore and Mouritsen, 2016; Xu et al., 2021; Wong et al., 2021). The well-studied Bogong moth, a nocturnal migrant, relies on both star (Dreyer et al., 2025) and magnetic (with complex connection to visual landmarks) (Dreyer et al., 2018a) information. However, knowledge from a single species cannot be generalized to other Lepidopteran migrants, especially because the multi-generational migrations of monarch butterflies and Bogong moths could be exceptional cases of highly adapted insect navigators. In both species, the same generation undertakes bidirectional movements (in autumn and spring) between breeding areas and specific geographic destinations - central Mexico for hibernation in monarchs, and the Australian Alps for aestivation in Bogong moths (Urquhart and Urquhart, 1978; Reppert and de Roode, 2018; Green et al., 2021). In most other Lepidopteran migrants, however, different generations perform multi-generational migrations between broad latitudinal breeding and overwintering ranges, rather than completing directed movements to a specific geographic location (Chapman et al., 2008; Chapman et al., 2015; Tong et al., 2022). Because most migratory insects travel between broad regions rather than pinpoint destinations, they likely experience only weak evolutionary pressure to develop highly accurate or complex orientation systems. Studies conducted on European common migrants, such as the painted lady butterflies *Vanessa cardi* and red admirals *Vanessa atalanta*, revealed that these species rely on a sun compass and but showing no evidence of a magnetic sense (Brattström, 2007; Nesbit et al., 2009; Pakhomov et al., 2025).

Little is known about the migratory behaviour and orientation mechanisms of other moth species. In Europe, several studies suggest that some species may use a magnetic compass (Baker and Mather, 1982) or moon/stellar cues (Sotthibandhu and Baker, 1979) for orientation and may even integrate information from the moon’s azimuth with the magnetic field (Baker, 1987). However, because these earlier studies were conducted on the British Isles without the use of modern flight simulators (e.g. Mouritsen and Frost, 2002; Dreyer et al., 2021), their findings are not easily comparable to more recent research in controlled environments. In Asia, the recently invasive fall armyworm (*Spodoptera frugiperda*), a dangerous pest species which has developed new migratory routes across China (Chen et al., 2023), relies on both geomagnetic and visual cues for accurate migratory orientation, resembling the mechanisms described in the Bogong moth (Ma et al., 2025). Further comparative studies across moth species with diverse migratory strategies are required to assess whether the orientation mechanisms observed in Australian Bogong moths represent a common feature among nocturnal Lepidopteran migrants. This knowledge gap served as the inspiration for the compass-focused part of the present study.

Despite the steadily increasing knowledge about orientation mechanisms in migratory moths, navigation (position-finding) abilities of all Lepidopteran migrants remain largely unexplored except for the monarch butterfly (Mouritsen et al. 2013a,b). To assess these abilities, researchers commonly conduct displacement experiments, which can be either physical or virtual. Most physical displacement studies on birds, in which animals are actually transported hundreds or even thousands of kilometres away from their typical migratory routes, have shown that adult migratory birds are able to compensate for such displacements by changing their flight direction or activity in orientation cages to compensate their displacement (Perdeck, 1958; Perdeck, 1967; Thorup and Rabøl, 2007; Chernetsov et al., 2008). In contrast, naïve (first-year) conspecifics usually rely on a simple clock-and-compass mechanism to reach their migratory destination (Mouritsen and Næsbye Larsen, 1998; Mouritsen, 1998b; Mouritsen and Mouritsen, 2000; Chernetsov and Utvenko, 2025; but see Thorup et al., 2020). Using virtual magnetic displacement experiments - manipulations when animals are exposed to geomagnetic conditions occurring at other locations on Earth without physically transporting them to those sites (Lohmann et al., 2022; Schneider et al., 2023) - researchers have shown that migratory birds (Kishkinev et al., 2015; Packmor et al., 2024), sea turtle hatchlings (Lohmann and Lohmann, 1996; Goforth et al., 2025) and adult sea turtles (Lohmann et al., 2004), salmon (Putman et al., 2014; Putman et al., 2020), European eel (Naisbett-Jones et al., 2017), sharks (Keller et al., 2021) and even newts (Fischer et al., 2001; Diego-Rasilla and Phillips, 2021) can detect and respond appropriately to such manipulations by compensating for the simulated displacements based on geomagnetic cues. Moreover, migratory birds can use one of the geomagnetic field (GMF) parameters - magnetic declination (the angle between true north (geographic north) and magnetic north (the direction a compass needle points)) - to determine longitude (Chernetsov et al., 2017), and are able to transfer magnetic map information from magnetoreceptors in the upper beak to the brain via the trigeminal nerve (Kishkinev et al. 2013; Pakhomov et al., 2018). They can navigate using geomagnetic cues extrapolated beyond their previous experience (Kishkinev et al., 2021) and rely only on two of the three GMF parameters - inclination (the angle between the Earth’s magnetic field lines and the horizontal plane at a given location) and declination - to extract positional information (Packmor et al., 2024). Whether migratory moths possess an ability to navigate using GMF similar to other animals mentioned remains underexplored question. This knowledge gap served as the inspiration for the magnetic displacement part of this present study.

To date, only one study has examined the navigational abilities of Lepidopteran migrants. In this experiment, North American monarch butterflies were physically displaced by 2,500 km from the eastern part of the United States to the west, yet their orientation in flight simulators remained unchanged compared with their pre-displacement orientation (Mouritsen et al., 2013a), suggesting that monarchs use a simple vector-navigation strategy (steering in a constant direction for a specific time or distance) without an ability to compensate for displacement (Mouritsen et al., 2013b; Oberhauser et al., 2013). A recent study on the same species examined the behaviour of migratory monarch butterflies under different magnetic conditions near their wintering grounds in Mexico, using a righting apparatus (Kendzel et al., 2023) and virtual magnetic manipulation and found no evidence of a magnetic map in this species (Shively-Moore et al., 2025), but its findings should be independently replicated using classical orientation methods such as flight simulators – a “gold standard” technique in studying orientation and navigation in Lepidopteran species (Mouritsen and Frost, 2002; Dreyer et al., 2021).

Here, we present the results of our study on migratory red underwings (*Catocala nupta*), previously tested in flight simulators at the same site in Austria (Dreyer et al., 2018b), to identify natural cues this species use for orientation and navigation. We conducted virtual magnetic displacement experiments, displacing this migratory moth species outside its natural European distribution range for the first time to investigate whether moths can use magnetic cues to compensate for such a displacement (magnetic map) extrapolating GMF beyond their species range. In addition, we tested red underwings in the absence of specific orientation cues, either magnetic (under a vertical magnetic field) or celestial (under simulated overcast conditions), to determine whether this species employs stellar and/or magnetic compasses (or both) to maintain its population-specific migratory direction during nocturnal migration.

After the virtual magnetic displacement, we expected several possible behavioural responses to this magnetic manipulation, based on findings from similar studies conducted on other migratory animals. If animals can perceive map information from the Earth’s magnetic field and use it to determine their position relative to their final destination, then being virtually displaced to a magnetically unfamiliar location (one they have never experienced before) should elicit a compensatory reorientation towards their migratory corridor. Such behaviour has been demonstrated in adult Eurasian reed warblers *(Acrocephalus scirpaceus,* hereafter reed warblers), which were able to compensate for virtual magnetic displacements by reorienting appropriately (Kishkinev et al., 2015; Chernetsov et al., 2017). Remarkably, this species can do so even when virtually displaced outside its natural distribution range (into areas where no conspecifics have ever been) by navigating using geomagnetic cues (inclination and declination) extrapolated beyond their previous experience (Kishkinev et al., 2021; Packmor et al., 2024). Similar compensatory reaction towards migratory corridors has been found in other migratory animals, such as sea turtles (Lohmann et al., 2004), salmons (Putman et al., 2020) and eels (Naisbett-Jones et al., 2017). Another possible response to such displacements - when animals can perceive navigational information from the geomagnetic field but are unable to interpret or use it correctly - is disorientation. This reaction has been demonstrated in bonnethead sharks (*Sphyrna tiburo*), which are able to compensate correctly for a virtual southward displacement from their capture site, recognising that a weaker magnetic field corresponds to more southerly locations. However, when exposed to a stronger magnetic field simulating a northward displacement (one they have never experienced), the sharks exhibited orientation, indistinguishable from random (Keller et al., 2021). A similar reaction has been observed in hatchling loggerhead sea turtles (*Caretta caretta*), which failed to orient in magnetic fields far outside their normal migratory route but were strongly oriented within the typical population range (Fuxjager et al., 2011). Comparable results were also found in juvenile reed warblers following a virtual westward displacement in a magnetic declination study (Chernetsov et al., 2017). Finally, if animals are unable to use all—or specific—parameters of the geomagnetic field and therefore lack a functional magnetic map, they cannot detect that they have been displaced hundreds or even thousands of kilometres from the capture site. In this case, they would be expected to orient in the same direction as they would under the magnetic field of their original location, like juvenile and adult European robins (*Erithacus rubecula*) and adult garden warblers (*Sylvia borin*) in a magnetic declination study (Chernetsov et al., 2020).

## Materials and Methods

### Moths’ capture and husbandry

Red underwings of both sexes were captured using a sugar-bait method, in which a cotton rope soaked with a mixture of red wine, sugar, honey, and bitters was wrapped around a tree (Sussenbach and Fiedler, 1999; Lucci Freitas et al., 2014) at the Biological Station Lake Neusiedl in Illmitz, Austria (47° 46’ 08.9”N, 16° 45’ 57.2”E) between 9 and 27 August 2025. This technique proved more effective for this species than conventional light traps, based on comparisons with different trap designs deployed at similar sites and advice from entomologists.

Upon capture, all moths were housed indoors in standard mesh cages (30 × 30 × 30 cm) placed in a sheltered place (in a laboratory with windows which exposed them to the natural local photoperiod). Some of the mesh cages were equipped with a portable locomotor activity monitor (pLAM; Raspberry Pi 3 with a night-vision IR camera) to record and analyse the activity patterns of the captured moths (Sondhi et al., 2022). The moths were kept under controlled conditions, including a constant temperature (22–24 °C) and 74-75% relative humidity. All captured moths were provided with a 10% honey solution which was renewed daily and maintained under these conditions for 1–3 days prior to testing, allowing them to recover from any capture-related stress. Orientation experiments were conducted in a large open area free of trees, located at a rural site approximately 5 km from the nearest village. The site had previously been used for magnetic orientation studies in birds due to its low levels of background electromagnetic noise and minimal light pollution, which do not interfere with avian magnetic orientation (Packmor et al., 2021; Packmor et al., 2024). All moths captured on a given night were randomly assigned to groups tested under different experimental conditions (see below) and were never tested under more than one condition, except for the control and displacement conditions. After orientation tests, all red underwings were released in good body condition (no wings damage) well before the migration of their conspecifics was finished.

### Flight simulator

All behavioural experiments were performed in modified version of a Mouritsen-Frost flight simulator which allow to obtain flight tracks and analyse orientation of tethered insects (Mouritsen and Frost, 2002; Dreyer et al., 2021; Dreyer et al., 2025). The flight simulator was made of non-magnetic materials (PVC, aluminium, PETG) and comprised three main components (Figure S1):

1. **a flight chamber**: a white plastic cylinder (diameter 45 cm, height 50 cm) placed vertically on a plastic/aluminium table. Inner wall of the cylinder was covered with a black curtain to create homogenous non-visual cues environment and eliminate reflections and thus any light artefacts from the optic flow we projected underneath the moths. The moth attachment apparatus consisted of a clear acrylic bar mounted on top of the cylinder, with an optical encoder positioned at its center. The 11-cm-long fine tungsten rod (0.5 mm in diameter) was connected to the encoder, serving as the encoder shaft. The edges of the white plastic cylinder limited a moth’s visual access to the sky to about 120°. A small plastic tube (20 mm in length, 2 mm in diameter, with a 0.5 mm hole) was attached to the distal end of the rod, enabling a moth to be quickly and securely connected to the rod.
2. **direction recording system:** A miniature optical encoder (US Digital, Vancouver, USA) was used to continuously record the rotations of the tungsten rod carrying a tethered moth (0–360°). Within the simulator, moths could freely rotate in the horizontal plane, and their azimuth relative to magnetic North was measured with a resolution of 3° every 200 ms. The encoder was connected to a laptop via a data acquisition (DAQ) device, and the data were stored as angle–timestamp pairs in a .txt file. The laptop was placed inside a custom-made box (5 side-closed), and its display was covered with a red gel filter to minimise stray light.
3. **optic flow system:** According to (Dreyer et al., 2021), dim and slowly moving optic flow projected beneath the moth (always moving from head to tail regardless of the moth’s orientation in the arena) provides additional motivation for sustained flight, although this effect may not be consistent across species (Ma et al., 2025). To generate such optic flow, we used a portable projector (Kodak Luma 75) enclosed in a 3D-printed plastic box with air vents to minimise stray light. The projector beam was directed via a 45° mirror onto a diffusing screen (Lee Filters 250 half-white diffuse) positioned beneath the arena (Figure S1). The optic flow intensity was reduced to nocturnal level (average luminance: 12×10^-4^ cd m^-2^ or 4×10^13^ photons m^-2^ s^-1^) using a combination of two ND0.9 neutral density filters (Lee Filters) and a variable circular ND 2–400 filter (KF Concept) placed over the projector’s exit aperture. The projector was connected to the laptop via a 4m HDMI cable, and custom-written software controlling the direction of image movement was integrated with the optical encoder system through a feedback loop (Figure S1B). This setup ensured that the optic flow consistently moved 180° relative to the moth’s heading (from head to tail) throughout the entire test.

### Test preparations and common procedures for all experimental conditions

Prior to attachment to the tethering stalks, each red underwing was kept in a fridge for 10 min to immobilise it. A small (3 cm length, 0.5 mm diameter) section of tungsten rod (ferromagnetic free) with footplate attached to a small piece of paper (a tethering stalk hereafter) was vertically glued to the moths’ thorax after all the scales were removed from the thorax. After this procedure, the red underwings were returned to a mesh cage, where they were kept for the rest of the day before being tested in the flight simulator. Before the experiments, mesh cages containing tethered moths were placed outdoors 30 minutes before local sunset to provide the animals with an unobstructed view of the sky, the setting sun, and the stars after sunset, in case these cues were important for compass calibration (Cochran et al., 2004; Pakhomov and Chernetsov, 2020).

All flight simulator tests were conducted between the end of nautical twilight - when no solar cues remain and the stars are clearly visible in the night sky - until 1:30 a.m. This period corresponds to the time when moths are typically active in the mesh cages (Figure S2). Experiments were performed only on moonless nights with clear skies (cloud cover below 20%) and under windless conditions. Temperature and wind speed were monitored throughout each test night using a portable weather station (Kestrel 5500, Nielsen-Kellerman Company, USA). During testing nights, the mesh cages containing the moths were placed inside cardboard boxes with an open top to prevent visual access to the local surroundings while providing an unobstructed view of the starry sky at the experimental site. When the local temperature dropped below 13°C, a small bottle filled with hot water and wrapped in a towel was placed inside the box to gently warm the moths before testing. Each moth was carefully removed from the mesh cage by grasping the stalk attached to its thorax. The stalk was then inserted into a small plastic tube attached to the rod in the flight simulator.

Each test lasted 5 minutes and was included in the further analysis only if 1) the moth was active continuously for the full 5 min (no more than 4 stops during tests), 2) it was flapping was vigorous, with a large and symmetrical amplitude for both wings, indicating that the stalk was securely attached to the moth’s thorax and did not interfere with its flight behaviour, 3) no continuous circling (spiraled) behaviour occurred, defined as persistent circular flight without pausing or directional choice during first 2 minutes after it was attached to the encoder. To minimize handling bias, the first two minutes of each flight were excluded from both recording and orientation analysis. The flight chamber was randomly rotated between trials, and the positions of the laptop and projector were changed each testing night to eliminate potential directional biases associated with the apparatus. Only head torches (headlamps) with weak red light were used to minimise potential effects of white light on the behavioural and physiological state of the tested moths.

### Virtual magnetic displacement experiments

For manipulating the magnetic field during virtual magnetic displacements, we used a three-axis, single-wound 2 m × 2 m × 2 m 3D Helmholtz coil system (the magnetic coils hereafter, Figure 1A, S1). The coil was identical to that used in previous virtual displacement experiments with songbird migrants (Kishkinev et al., 2021; Packmor et al., 2024) and magnetic orientation experiments with migratory monarch butterflies (Mouritsen and Frost, 2002), so full details of the specifications can be found in these references. This coil system generates a magnetic field with >99% homogeneity of the applied field strength (<200 nT) in the central area where a flight simulator and a cage with moths were placed. Each of the three axes of the coils was powered by a separate power supply BOP 50-4 M (Kepco Inc., USA), allowing precise control of the field’s intensity and direction. The parameters of the magnetic field were checked in the centre of the flight simulator and the cage using fluxgate 3-Axis MR3 Magnetoresistive Milligauss Meter (AlphaLab Inc., Salt Lake City, Utah, USA) with a resolution of 1 nT.

**Figure 1.**
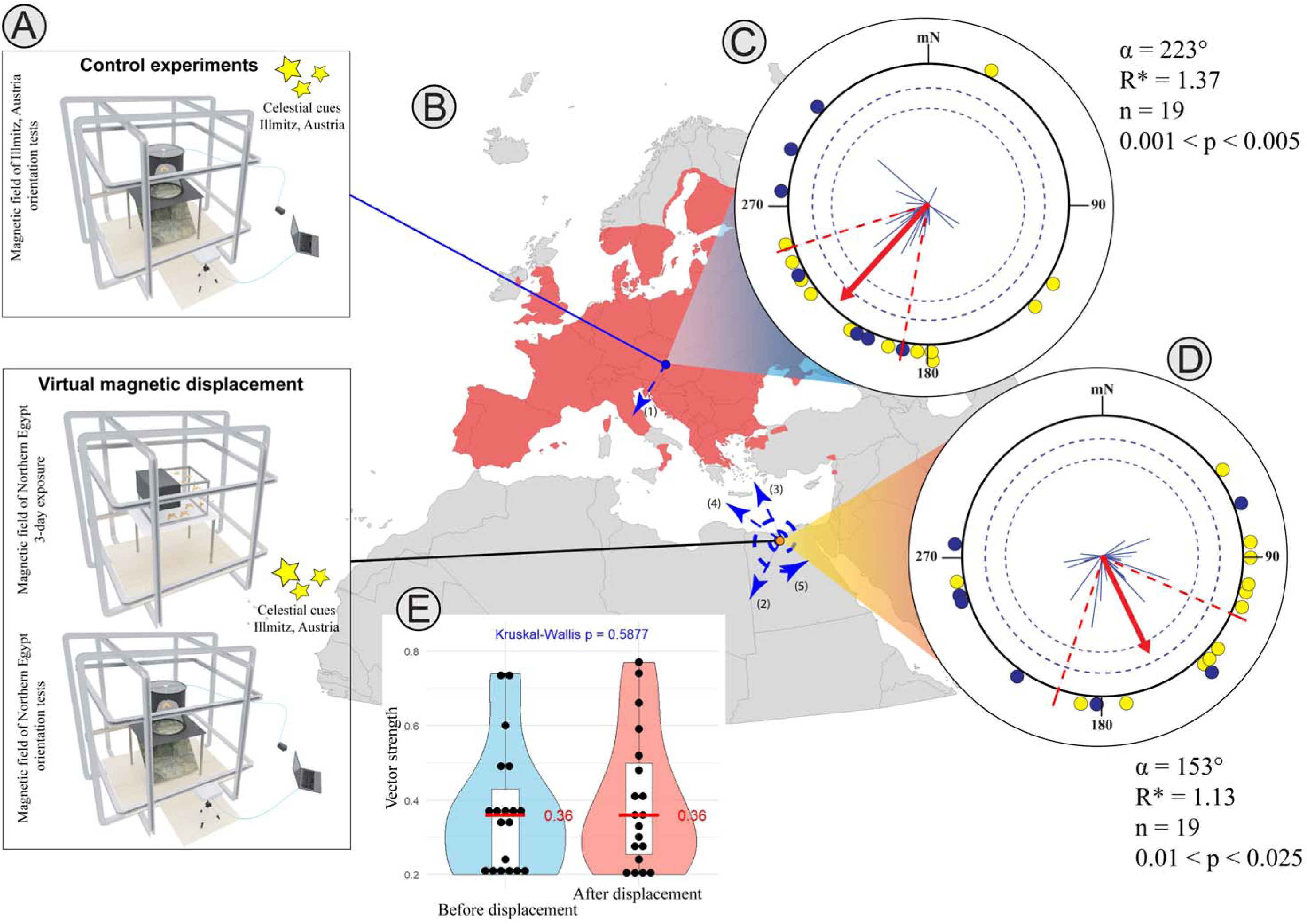
Virtual magnetic displacement experiments. **A:** schematic illustration of the experimental setup and experimental conditions; **B**: Distribution of red underwings in Europe, based on data from iNaturalist, Obsidentify, GBIF (Figure S3) and (Leraut, 2019); **C**: Orientation of the moths before displacements (in the magnetic field of Austria); **D**) orientation of the moths after the virtual displacement (to the magnetic field of Northern Egypt): **E**: Directedness of the individually tested moths in before and after virtual displacement (vector strength, r values, black dots). Each dot at the circle periphery indicates the mean orientation of one individual moth (yellow – first tested group, blue – second tested group). Dashed lines indicate the significance thresholds of the MMRT for 5% and 1%, respectively. The mean direction of the red underwing moths is shown as a red line, blue lines – individual vectors of tested moths, mN – magnetic North. The red radial dashes show the 95% confidence intervals.

Before virtual magnetic displacements, all red underwings were tested under control conditions with access to celestial cues (stars and Milky Way) and in the unchanged, natural magnetic field (NMF: total magnetic intensity∼49140 nT, magnetic inclination∼64.9°, magnetic declination∼4.7°, Figure 1A). The flight simulator was placed within magnetic coils (Figure S1C) that were not of the double-wrapped Merritt design typically used in our previous studies and allowing current to flow in two directions (Kishkinev et al., 2015; Pakhomov et al., 2018). In a double-wrapped coil system, when the current flows in the parallel direction, an artificial magnetic field is generated, whereas an antiparallel current cancel the artificial field, leaving the resultant field identical to the natural geomagnetic field (ideal control condition). In the present setup, it was not possible to run current in antiparallel directions to reproduce the normal magnetic field (NMF) while the coils were active. Therefore, to minimise potential effects of noise from the power supplies on the results of subsequent virtual magnetic displacement experiments, the power supplies were switched on during control tests but not connected to the coils similar to previous study on songbird migrants (Packmor et al., 2024) and they produced the same amount of noise in all conditions to control for any potential noise effects on the moths’ behaviour.

After the control tests, the flight chamber of the simulator was removed from the magnetic coils and all moths that met the inclusion criteria described above (first group: n = 12; second group: n = 7) were transferred to a custom-made non-magnetic mesh cage (CMF exposure cage, ∼50 × 40 × 40 cm) placed on the flight simulator table (Figure S1D). This cage allowed the moths to see the local surroundings, the sun at sunrise and sunset, and a clear starry sky. It also included a shelter (an area with shade under a partially covering cage roof) where they spent the daytime, which likely reduced stress associated with prolonged outdoor exposure. Using the magnetic coils, we generated a new magnetic condition simulating the geomagnetic parameters of northern Egypt (near Alexandria: CMF total intensity ∼ 44,300 nT; inclination ∼ 46°; declination ∼ 4.8°, Figure 1B, S4). These parameters do not create multiple simulated locations and correspond to a single area that includes northern Egypt, according to the ViDMAL tool (Figure S4, Schneider et al., 2023). The magnetic field parameters for the virtual displacement were calculated using the NOAA online magnetic field calculator based on the 2025 World Magnetic Model (WMM). Because the exact wintering site of this population of red underwings is unknown, we chose to virtually displace them outside their typical European distribution range (Figure 1B, 3S) - following the design of previous virtual magnetic displacement experiments on reed warblers (*Acrocephalus scirpaceus*) conducted at the same site (Kishkinev et al., 2021). These earlier studies demonstrated that these birds could compensate for such displacements beyond their natural range and extrapolate from unfamiliar magnetic parameters (Kishkinev et al. 2021; Packmor et al 2024).

The moths were kept in the changed magnetic field (CMF) for three days, fed daily with a fresh 10% honey solution, and attached to tungsten stalks without leaving the central part of the coils, where the CMF was >99% homogeneous (<1% variation in the amount the magnetic field was changed, which means that in the experiments reported here, <50 nT variation in the field was caused by the coils as the field was changed just under 5,000 nT). To this, the natural daily variation in the field of typically 30-100 nT has to be added to get to the complete field variation range). After the three-day exposure to the simulated magnetic field of northern Egypt, the moths were transferred (30 minutes before the end of nautical twilight) to the same mesh cage (placed inside a cardboard box) used in the control experiments. The CMF exposure cage was then removed from the magnetic coils, and the flight chamber was placed back on the table for the orientation tests conducted later that night. The cardboard box with the mesh cage was kept inside the central part of coils with homogeneous magnetic field during the orientation tests. To sum up, the CMF moths never left the homogenous CMF area of the coil during keeping and testing during the CMF experimental period. Fine adjustments and regular calibrations of the magnetic field were performed before and after testing each group of moths to ensure that the desired magnetic conditions were accurately maintained at the centre of the setup. Two groups of red underwings were tested under both control conditions and during virtual magnetic displacement experiments:

1. First group (n = 12): control tests conducted on 12–15 August; tests following virtual displacement conducted on 23 August.
2. Second group (n = 7): control tests conducted on 26–27 August; tests following virtual displacement conducted on 31 August.

### Experiments under overcast and VMF conditions

In addition to virtual magnetic displacement experiments, another group of moths (not subjected to displacement) was tested under simulated overcast conditions and in a vertical magnetic field (VMF) to assess whether red underwings rely on geomagnetic or celestial cues to maintain their population-specific migratory direction. All orientation experiments under such conditions were conducted in the flight simulator placed inside the magnetic coils mentioned above. Under simulated overcast conditions, the upper part of the flight simulator was covered with a UV-transmissive diffuser (Lee 251 Quarter White gel filter; ∼80% light transmission, no spectral distortion), eliminating visual access to the starry sky while preserving the natural spectral composition and intensity of ambient light, ensuring that the geomagnetic field was the only available compass cue. This approach is commonly used to study magnetic orientation in the absence of visual cues in other animals, including migratory birds and butterflies (Mouritsen and Frost, 2002; Bojarinova et al., 2023). In the NMF control condition, the power supplies of the magnetic coils were turned on, but were not connected to the coils, thus creating the same level of acoustic noise as during control and VMF experiments. Tests under simulated overcast conditions were conducted on 13 August - 01 September.

To investigate whether red underwings use a stellar compass, we tested them in the same flight simulator with an open top, allowing access to a clear starry sky. The simulator was placed inside magnetic coils that generated a vertical magnetic field (VMF) of normal Earth intensity (NMF = 49120 ± 39 nT; VMF = 49093 ± 24 nT), which provided the tested moths with no usable magnetic compass information due to absence of the horizontal components or its small value (VMF inclination: 89.5° ± 0.4° SD). It has been indicated that night-migratory songbirds cannot use a magnetic field for their compass if the inclination is steeper than 88° (Lefeldt et al., 2015) and the vertical magnetic field (VMF) manipulation is highly effective for disrupting magnetic orientation, as demonstrated in previous star compass studies on migratory birds (Pakhomov et al., 2017; Zolotareva et al., 2021) alongside near-zero or true zero magnetic field (ZMF) conditions (Mouritsen, 1998; Dreyer et al., 2025). The VMF may be more suitable for behavioural experiments, as it better mimics natural conditions (maintaining the same field intensity as the normal magnetic field, NMF) and no studies have reported any negative effects of this manipulation on behavioural or physiological traits of animal, unlike ZMF (Wan et al., 2020; Krylov et al., 2022; Bar-Ziv et al., 2025, but see Dreyer et al., 2025). Tests under VMF were conducted on 25 August - 01 September.

## Statistical analysis

The classical Rayleigh test of uniformity (RT) was used to assess whether the group mean orientation differed significantly from the uniform distribution (Batschelet, 1981). In addition, second-order statistics were conducted using the nonparametric Moore’s Modified Rayleigh Test (MMRT), which weights individual mean angles according to their r-values (Moore, 1980) implemented via a modified version of the R script from (Massy et al., 2021). For both tests (RT and MMRT), a low p-value (p < 0.05) indicates that the tested moths oriented in a certain direction and that their orientation data were significantly different from a uniform distribution. All individual orientation experiments with r < 0.2 were excluded from the analysis, as such trials were assumed to reflect behaviours different from migratory orientation according to previous studies on butterflies and moths (Nesbit et al., 2009; Dreyer et al., 2018b).

Differences in orientation between different experimental groups of red underwings were evaluated using the nonparametric Mardia–Watson–Wheeler test (MWW), and the Moore’s paired test was applied for comparisons of the same individuals tested under different conditions. To further characterise orientation patterns under various experimental treatments, we conducted maximum-likelihood analyses using the CircMLE R package (Fitak & Johnsen, 2017). Differences in r-value between the moths from different groups was analysed using the non-parametric Kruskal-Wallis test.

Additionally, we applied a bootstrap technique to test whether significantly oriented groups showed higher directional consistency than disoriented ones (Leberecht et al., 2022; Romanova et al., 2023). In this approach, a random sample of orientation directions (n angles) was drawn with replacement from the individual moth orientation directions in the significantly oriented group. For each resample, the corresponding r-value was calculated, and this procedure was repeated 100,000 times. The resulting r-values were ranked in ascending order, and the values at ranks 2,500 and 97,500 (and at 500 and 99,500) defined the 95% and 99% confidence limits for the r-value of the significantly oriented group, respectively. If the observed r-value of the disoriented group was lower than these intervals, the oriented group was considered significantly more directed than the disoriented one at p < 0.05 or p < 0.01, respectively. Bootstrap and maximum-likelihood analyses, as well as MMRT and Kruskal-Wallis test, were conducted in R version 4.3.2. (The R Core Team, 2013). The MWW test was performed in Oriana 4.02 (Kovach Computing Services, UK), while RT calculations and RT orientation visualizations for each group were performed using a custom-written Python script.

## Results

### Virtual magnetic displacement

Red underwing moths were significantly oriented and preferred a southwestern direction in flight simulator experiments (RT: α = 217°, r = 0.61, n = 19, p < 0.001, 95% CI = 191° – 245°; MMRT: α = 223°, R* = 1.37, n = 19, 0.001 <p < 0.005, 95% CI = 190° – 253°, Figure 1C; see individual moth headings in Table S1 and CircMLE modelling results in Table S2A) when they had full access to the natural starry sky along with an undisturbed natural magnetic field of Illmitz, Austria, i.e. before any virtual displacements. After the virtual magnetic displacement, several possible orientation responses of red underwings to this magnetic manipulation could be expected:

1. No reaction to the displacement and continuous flying in their typical south-western migratory direction (Figure 1B(2));
2. Detection of the displacement as being outside their normal migratory route and attempt to return to the study site in Austria, orienting towards the north-north-west (Figure 1B(3));
3. Recognising the displacement and attempt to compensate for it by orienting towards the theoretical wintering area in a west-north-west direction (e.g. Spain or Italy; Figure 1B(4)); or
4. Detection of the displacement as inconsistent with their expected route, resulting in disorientation (Figure 1B(5)).

However, after three days of exposure to Egypt’s magnetic field, the preferred mean direction of the same red underwings was significantly oriented towards a somewhat unexpected south-southeast (RT: α = 145°, r = 0.46, n = 19, p = 0.02, 95% CI = 112° – 195°; MMRT: α = 153°, R* = 1.13, n = 19, 0.01 <p < 0.025, 95% CI = 114° – 198°, Figure 1D; CircMLE modelling results: Table S2B). However, there were no significant differences in r-values before and after the virtual magnetic displacement (Kruskal–Wallis test: χ² = 0.294, p = 0.5877; Figure 1E), and the 95% confidence intervals of the mean directions slightly overlapped. Nevertheless, the difference between the two circular distributions obtained before and after the virtual magnetic displacement approached significance (Moore’s test: R′ = 1.21, p < 0.025) most probably due to the tendency to a difference in the mean group orientations (though formally shy of being statistically significantly different).

### Orientation of red underwings in absence of magnetic or celestial cues

When access to celestial and other visual cues was eliminated under simulated overcast conditions (Figure 2A), red underwings lost their ability to maintain the population-specific migratory direction observed under control conditions (RT: α = 200°, r = 0.25, n = 16, p = 0.35; MMRT: α = 231°, R* = 0.32, n = 16, 0.5 < p < 0.9, Figure 2B; CircMLE modelling results: Table S2C). Notably, under a natural night sky but in the absence of geomagnetic compass information (VMF), the moths oriented in their seasonally appropriate south-westward migratory direction (RT: α = 201°, r = 0.74, n = 13, p < 0.001, 95% CI = 176° – 226°; MMRT: α = 211°, R* = 1.55, n = 13, p < 0.001, 95% CI = 191° – 237° Figure 2C; CircMLE modelling results: Table S2D) and the orientation in VMF did not differ significantly (MWW test: W = 0.341, p = 0.84) from the orientation of red underwings under the control condition (NMF before displacement, Figure 1C).

**Figure 2.**
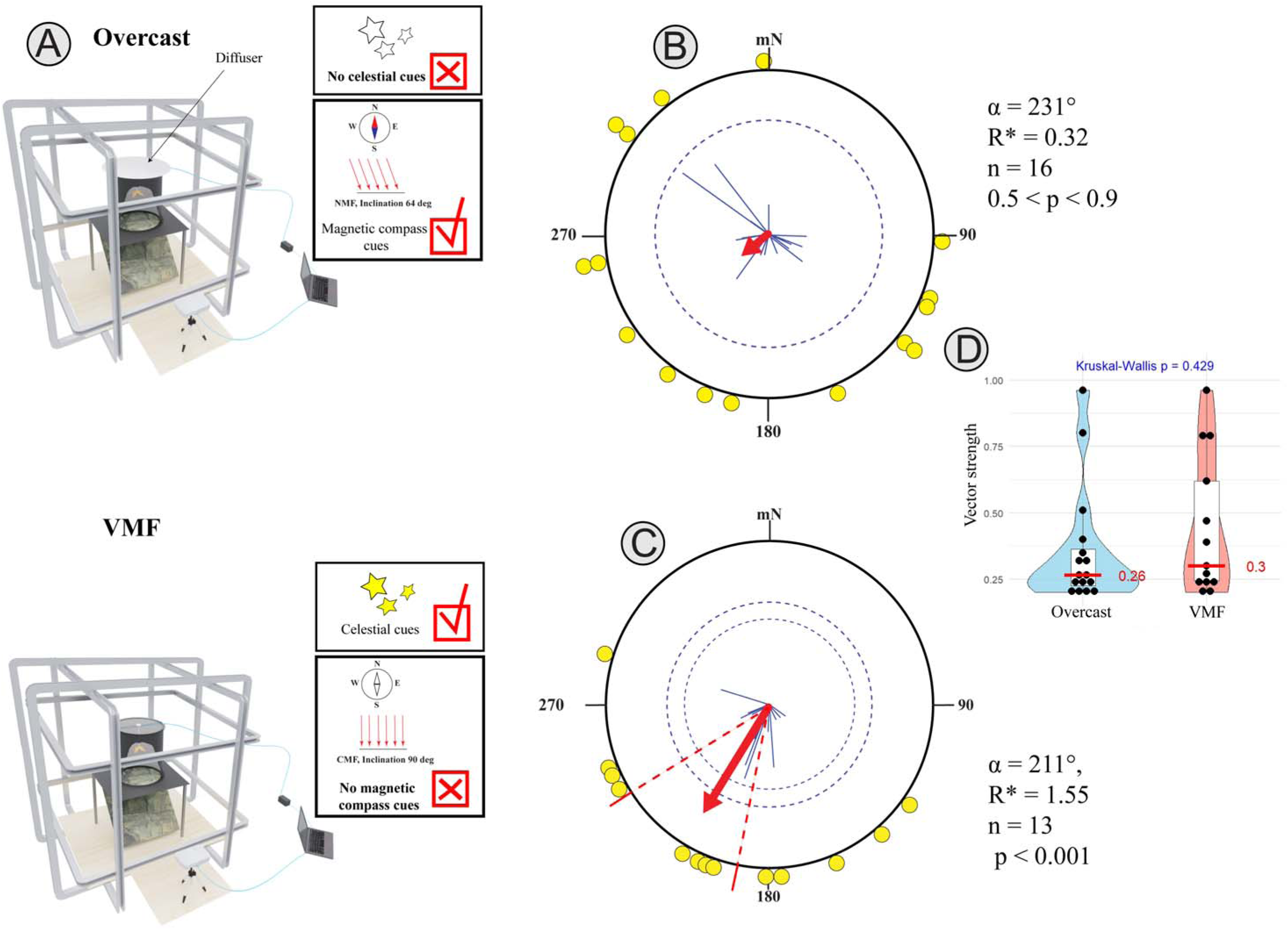
Orientation of red underwings with access only to magnetic or celestial cues. **A:** schematic illustration of the experimental setup and experimental conditions; **B:** orientation of red underwing moths under simulated overcast condition; **C**: orientation of red underwings in the vertical magnetic field (VMF); **D**: Directedness of the individually tested moths under overcast condition and VMF conditions (vector strength, r values, black dots). Other details, see legend in Figure 1.

Importantly, the absence of visual cues did not reduce the moths’ motivation to fly in the experimental setup, as indicated by the lack of difference in r-values between groups (Kruskal–Wallis test: χ² = 0.626, p = 0.429, Figure 2D). In contrast, the flight directions of the moths under overcast conditions were significantly more scattered than those of the control and VMF groups, according to bootstrap analyses (Control vs. Overcast: p < 0.01; 95% CI for r-value: 0.34–0.79; 99% CI: 0.27–0.85, Figure S5A; VMF vs. Overcast: p < 0.01; 95% CI: 0.61–0.88; 99% CI: 0.56–0.91, Figure S5B).

## Discussion

### Control direction of red underwings compared to past works

In Europe, red underwing moths are widely distributed, except in the northern parts of Scandinavia, and they are absent from most Mediterranean inland regions, occurring only on Corsica and the Balearic Islands (Figure 1B, S3; (Leraut, 2019). Over the past two decades, the migratory status of this species has shifted from being considered “occasional vagrants” (Skinner, 2009), invading new areas only during favourable seasons (Fox et al., 2011), to regular autumn migrants moving from central Europe towards the Mediterranean region (Dreyer et al., 2018b). Recent experiments conducted at the same study site in Austria as the present study suggested that red underwings display a strong south-easterly orientation in flight simulator experiments (Dreyer et al., 2018b). According to that study, a south-easterly directional preference suggests that individuals in the wild caught at the Biological Station Lake Neusiedl in Illmitz may migrate further into the Balkans. In contrast, in our study, red underwings tested in the flight simulator under a starry sky and natural magnetic field (NMF) conditions preferred a more southwestern direction (Figure 1C,D), similar to that observed in large yellow underwings (*Noctua pronuba*) under the similar experimental conditions at the same place (Dreyer et al., 2018b).

The difference in mean direction under control conditions between our experiments and the previous study (south-west vs. south-south-east, respectively) might be explained by population-level variation, with distinct red underwing populations migrating to different parts of the Mediterranean region and exhibiting corresponding differences in their orientation responses in flight simulators. Moths in the study by Dreyer et al. (2018b) were captured during the first half of September using light traps and a searchlight, which primarily attract high-flying individuals. In contrast, in our present study, moths were collected using sugar baits during the second half of August, as our attempts to capture them with light traps similar to those used in Dreyer et al. (2018b) were unsuccessful. These differences in capture method and timing suggest that the moths studied by Dreyer et al. (2018b) may have originated from more northern regions relative to Austria and were migrating towards the Balkans, whereas the moths in our study were likely more local (from other parts of Austria or neighbouring countries) and migrating towards, for example, Italy during autumn migration. Differences in the migratory direction of free-flying and tethered Lepidopteran migrants from different populations migrating between broad latitudinal zones are not uncommon. For example, migratory red admirals exhibit considerable variation in their mean orientation, with flight directions ranging broadly from east to south-west across their European migration range (see Figure 1 in Pakhomov et al., 2023).

### Virtual magnetic displacement in red underwings suggests the lack of magnetic navigation

The results of our virtual magnetic displacement study show no evidence of compensatory reorientation (i.e. correction towards the migratory route) in red underwings following the apparent virtual southward displacement, even though their orientation in the magnetic field of Egypt might have differed slightly (borderline significance) from that of the same individuals before the displacement. Our interpretation is based on the fact that the south-easterly direction does not correspond to reorientation towards either the capture site (NNW) or any likely wintering sites in Italy or Spain (WNW or W), but rather aligns more closely with the southward direction that this species would take in the wild, as if the virtual displacement had never occurred.

A possible, though speculative, explanation for the south-easterly orientation observed after the displacement is that the two groups of red underwings, tested on separate days following their control experiments, might have expressed different migratory programmes. Moths from the first group (n = 12), which were virtually displaced at least seven days after the control experiments (due to unsuitable weather conditions), were oriented towards the south-east, whereas individuals from the second group (n = 7), which were kept for less than five days between control tests and displacements, preferred a more south-westerly direction (see directions showed by these two groups in Table S2B). Results from the maximum-likelihood analysis indicate that the post-displacement orientation distribution was more bimodal than unimodal, with the modelled directions closely matching those observed in the two groups (Table S2B). Consequently, because little is known about the migratory phenology, behaviour, and wintering grounds of this nocturnal migrant, the observed differences in preferred direction between two groups can, for instance, be attributed to seasonal progression. In nocturnal migratory birds, first-year inexperienced individuals follow an innate spatiotemporal programme (known as the ‘clock-and-compass’ or ‘calendar-and-compass’ mechanism) to reach their wintering grounds during their first migration (Berthold 1991; Mouritsen 2003; Chernetsov and Utvenko, 2025). According to this concept, juveniles fly in one direction for a certain number of days (n ), then change direction and continue for another period (n ), and so on, eventually reaching their population-specific wintering grounds (Gwinner and Wiltschko, 1978; Mouritsen 1998b, 2003; Mouritsen & Mouritsen 2000). A similar programme may also exist in nocturnal moths if they use specific sites to spend the winter months. Such a programme could guide them to these regions using a simple vector-based navigation strategy, without the need for a sophisticated navigation system based on geomagnetic cues (i.e. a magnetic map). Further research is needed to investigate whether nocturnal moths have similar spatiotemporal program to that of young night-migratory songbirds that leads them towards wintering regions. However, both the southwestern and southeastern directions observed in the present study may represent the species’ typical natural migratory orientation in Austria. The mean direction after magnetic displacement (153°) is close to the 146° previously reported (Dreyer et al., 2018b), suggesting that the observed headings fall within the natural orientation range of red underwings tested in Illmitz (coarsely southbound). This explanation is therefore more plausible than an effect of magnetic displacement or hypothetical ‘calendar-and-compass’ mechanism in moths.

The absence of a compensatory response in red underwing moths to a long-distance virtual displacement based on magnetic cues only is consistent with previous real displacement studies on diurnal monarch butterflies, which showed that they rely on a vector-based navigation strategy even after a 2,500 km physical displacement (Mouritsen et al., 2013a). Furthermore, monarchs do not respond to simulated magnetic conditions mimicking regions around their wintering grounds in Mexico (Shively-Moore et al., 2025), and the monarchs’ overwintering locations and population abundance remain unaffected by natural shifts in the Earth’s magnetic field (Guerra et al., 2022). All of these findings suggest that, just as in naïve, young night-migratory songbirds on their first autumn migration, Lepidopteran migrants primarily rely on vector navigation based on visual and geomagnetic compass cues rather than being true navigators (only experienced night-migratory songbirds can use a magnetic or other type of map during migration, but only after having completed at least the outward journey once (Chernetsov and Utvenko, 2025). Juveniles, by contrast, do not yet have such navigational ability (but see Thorup et al., 2020). This framework may help explain why neither monarch butterflies nor red underwings show any evidence of true navigation (the ability to find direction leading to a goal from an unfamiliar location in absence of cues emanating from the goal): due to their multi-generational annual cycle, only inexperienced individuals undertake the migration from the breeding grounds to the wintering areas, similar to first-year songbirds. However, in other migratory animals such as sea turtles and fish, juvenile individuals possess an inherited magnetic map or at least sensible responses to various magnetic fields they encounter on their journeys, as long as they remain within the bounds of their natural migratory corridor (Lohmann et al., 2001; Lohmann and Putman, 2007; Fuxjager et al., 2011; Putman et al., 2013; Putman et al., 2014; Brothers and Lohmann, 2018). In relation to Lepidopteran migrants, the migrations of monarch butterflies and Bogong moths are unique because the *same* individuals undertake the movement from the breeding grounds to non-breeding sites, where they hibernate or aestivate, and then initiate the return migration several months later and this pattern resembles the migration cycle of songbirds. Therefore, individuals that perform the spring migration in monarchs or the autumn migration in Bogong moths may resemble avian experienced migrants and may be the best candidates for virtual magnetic displacement experiments in the future to test this possibility.

### Evidence for the use of stars but not the geomagnetic field only for maintaining migratory direction

A variety of celestial cues such as the moon’s position, star patterns, the Milky Way, moon-derived polarized light, and the geomagnetic field (Baker and Mather, 1982; Foster et al., 2018; Goforth and Merlin, 2025) has been suggested to be used by nocturnal insect migrants for orientation. However, until recently, very little was known about whether nocturnal long-distance migratory insects can use celestial or geomagnetic cues for orientation. Recent work on Bogong moths has shed important light on this question: these moths seem to rely on the geomagnetic field **only when it is available together with visual cues**, and only when a conflict is created between magnetic north and the direction indicated by visual cues, they seem to become disoriented (Dreyer et al., 2018a; Ma et al., 2025). Red underwings in our study were unable to maintain the appropriate population-specific seasonal direction and instead showed disorientation (based on all statistical analyses applied, including the Rayleigh test, Moore’s Modified Rayleigh test, maximum-likelihood estimation, and bootstrap procedures) when they had access only to the geomagnetic field (under simulated overcast condition). This suggests two possible explanations:

1. red underwings might not use geomagnetic compass information for orientation, or
2. they may require magnetic information to be transferred to or integrated with visual cues (local landmarks), as demonstrated for Bogong moths and fall armyworms.

Importantly, our flight simulator provided no visual orientation information on its walls: the insects were tested under a diffuser inside a uniform black cylinder, with no artificial landmarks and no celestial cues. Interestingly, magnetic orientation in another migratory noctuid, the fall armyworm from China, is strongly light-dependent and fails completely in darkness (Ma et al., 2025) as well as large yellow underwing moths tested in a dark-room or with painted eyes (Sotthibandhu and Baker, 1979). Therefore, further experiments manipulating visual landmarks, natural or artificial celestial cues, and tests under a range of light intensities and spectra will be essential to determine whether red underwing moths do not use magnetic cues or require specific visual–magnetic interactions to orient during migration.

When we created the opposite experimental condition in which the clear starry sky was the only available compass cue (i.e., under VMF, where no geomagnetic compass information was provided), red underwings flew in their seasonally appropriate migratory direction, similar to their behaviour when they had access to both celestial and geomagnetic cues (control). These results suggest that these noctuid migrants can use the distribution of stars or star-derived cues (star constellation, Milky Way, etc) in the night sky alone to determine their required migratory heading. Our results for the first time provide evidence that European migratory moths possess a star compass similar to that of night-migratory birds (Emlen, 1970) and Bogong moths (Dreyer et al., 2025). As in the previous study on red underwings and large yellow underwings (Dreyer et al., 2018b), the red underwings in our experiments did not show any systematic change in their bearing throughout the testing nights (Figure S6), whether tested with access to both the geomagnetic field and stars (yellow line) or with access to stars only (red line). This may indicate that the star compass of red underwings is time-compensated or that they have learned the location of the celestial center of rotation, as in the star compass of night-migratory birds, which is time-independent (Mouritsen, 1998; Pakhomov et al., 2017). However, to test any dependence of the star compass on time of the day, further experiments with clock shifted moths are required. This was beyond the scope of this study.

## Conclusion

Based on our results, European migratory moths, red underwings, seems to rely primarily on night-time celestial cues (presumably stellar ones) to determine their population-specific migratory direction, which may differ among populations. Identifying which celestial cues (e.g., star constellations, the Milky Way, or other features of the night sky) are used by this species or other European moths and determining whether their star compass is time-dependent or not, will require further orientation experiments using projected starry skies and clock-shift manipulations. We did not find any evidence of magnetic orientation when this species was tested in the absence of visual cues. However, based on the data we present here, we cannot conclude that red underwings do not use magnetic cues at all, because Australian Bogong moths and armyworms in China seem to need interactions between visual and magnetic information in order to use their magnetic compass. The results of our virtual magnetic displacement experiments clearly indicate that red underwings do not use parameters of the Earth’s magnetic field to determine their position or to compensate for displacements to unfamiliar places. Similar to North American monarch butterflies, our data on red underwing moths suggests that the moths rely on a simple vector-navigation strategy rather than true navigation to reach their wintering grounds (Mouritsen et al., 2013a; Warrant and Maleszka, 2026).

## Acknowledgments

We thank all members of the Biological Station Illmitz, in particular Justin Catau and Gilbert Hafner, for their assistance with logistics and permit procedures, and Nazar Shapoval for his advice on catching red underwings.

## Authors’ contributions

A.P.: conceptualization, data curation, formal analysis, funding acquisition, investigation, methodology, project administration, visualization, writing—original draft; D.K.: conceptualization, funding acquisition, methodology, writing—review and editing; D.D.: methodology, resources, writing—review and editing, T.Z.: methodology, resources, writing—review and editing; H.M.: resources, writing—review and editing.

## Data availability statement

All data (raw data of experiments, results of additional statistical analysis, etc) and Python/R scripts are provided in electronic supplementary material and posted on Zenodo (https://doi.org/10.5281/zenodo.18852923). All components of the flight simulator, Python and R scripts, 3D printing/laser cutting files can be found here: https://magbbb.com/openscience/.

## Ethical considerations

All applicable international, national and/or institutional guidelines for the care and use of animals were followed. The experiments were conducted in accordance with the national animal welfare legislation of Austria and with permission of the state of Burgenland (Abteilung 4 - Ländliche Entwicklung, Agrarwesen und Naturschutz; permit: A4-HAS-RNS-2025-008.662-1/4). Additionally, the experiments received local ethical approval by the animal welfare ethics review body (AWERB) of Keele University as the core research team (A.P. and D.K.) were employed by the organization during the period of data collection.

