## Supplemental tables and Figures for "The Role of Magnetic and Celestial Cues in Orientation and Navigation of Red Underwing (*Catocala nupta),* a European Migratory Moth"

**Figure S1. A flight simulator to study orientation and navigation in migratory moths**

**A)** (A) 3D model of the flight simulator integrated with magnetic coils, optic flow, and stellar sky systems (left), and its schematic diagram (right).

(B) Schematic of the upper part of the apparatus, showing the cylinder, optical encoder, and tethered insect, together with the feedback loop system for optic flow.

(C) Photograph of the flight simulator with the magnetic coil system.

(D) Photograph (left) and 3D model (right) of the cage positioned inside the magnetic coils during exposure to the magnetic field of northern Egypt.

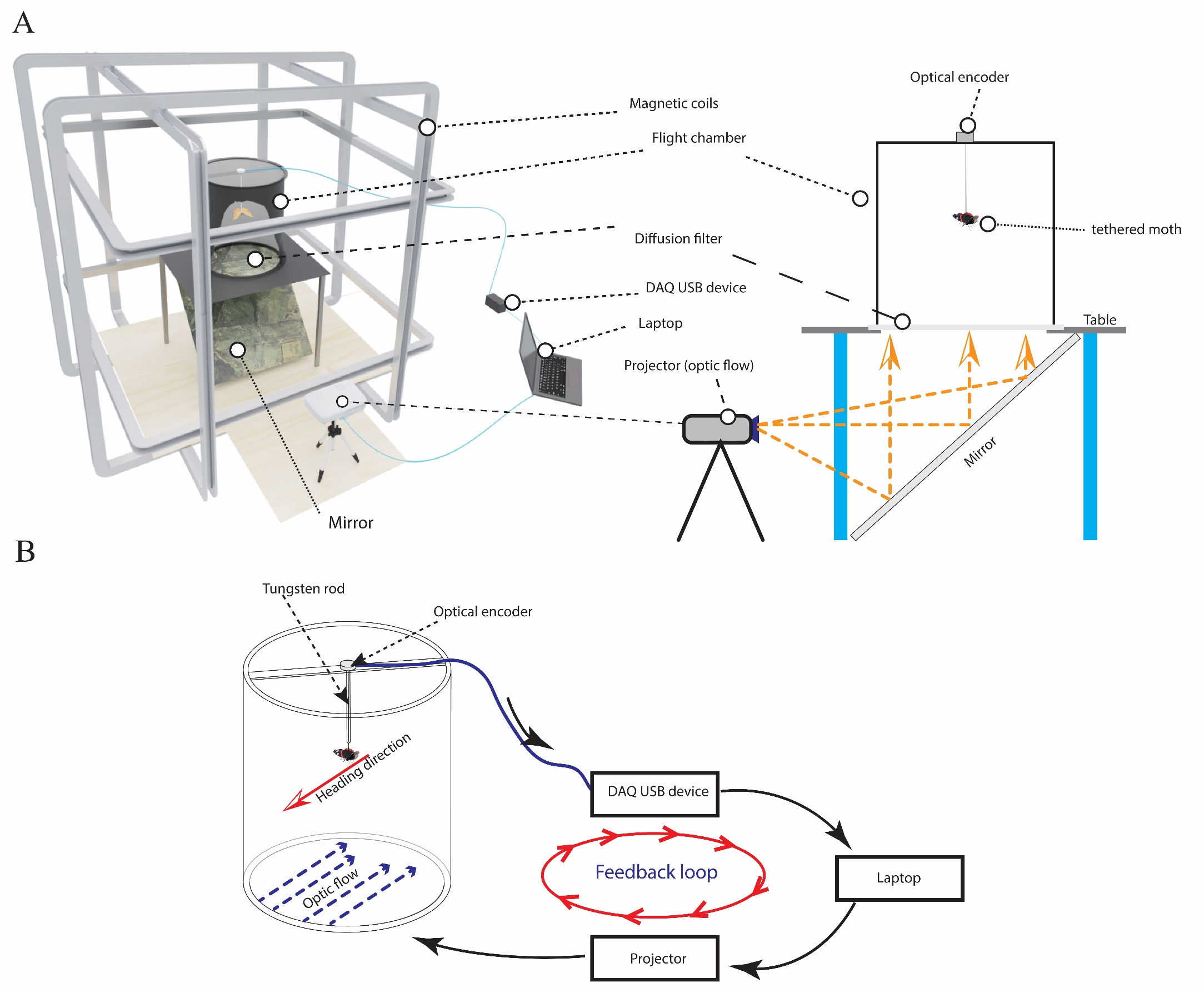

| **C** |  |  |
| --- | --- | --- |
| 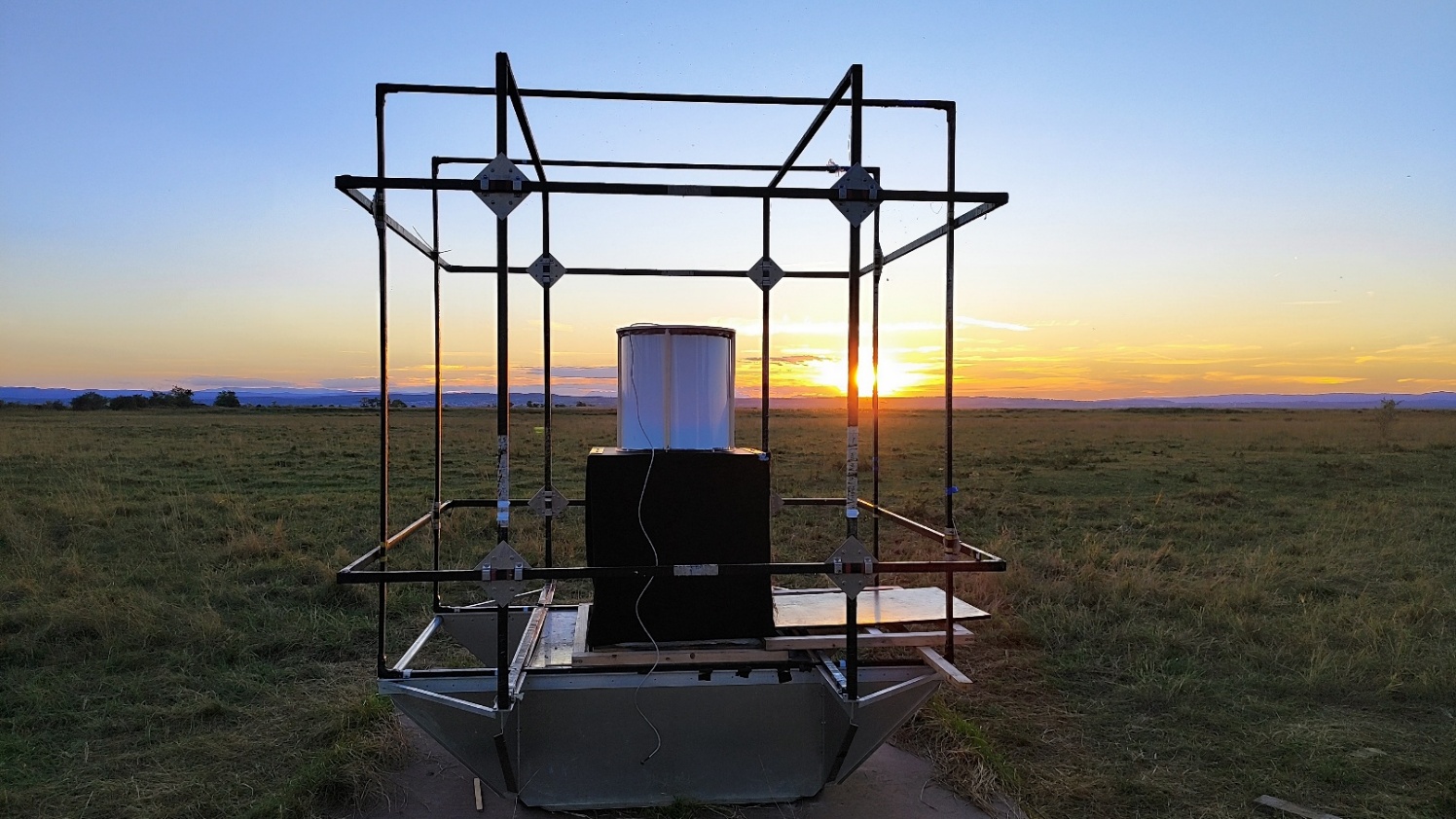 |  | 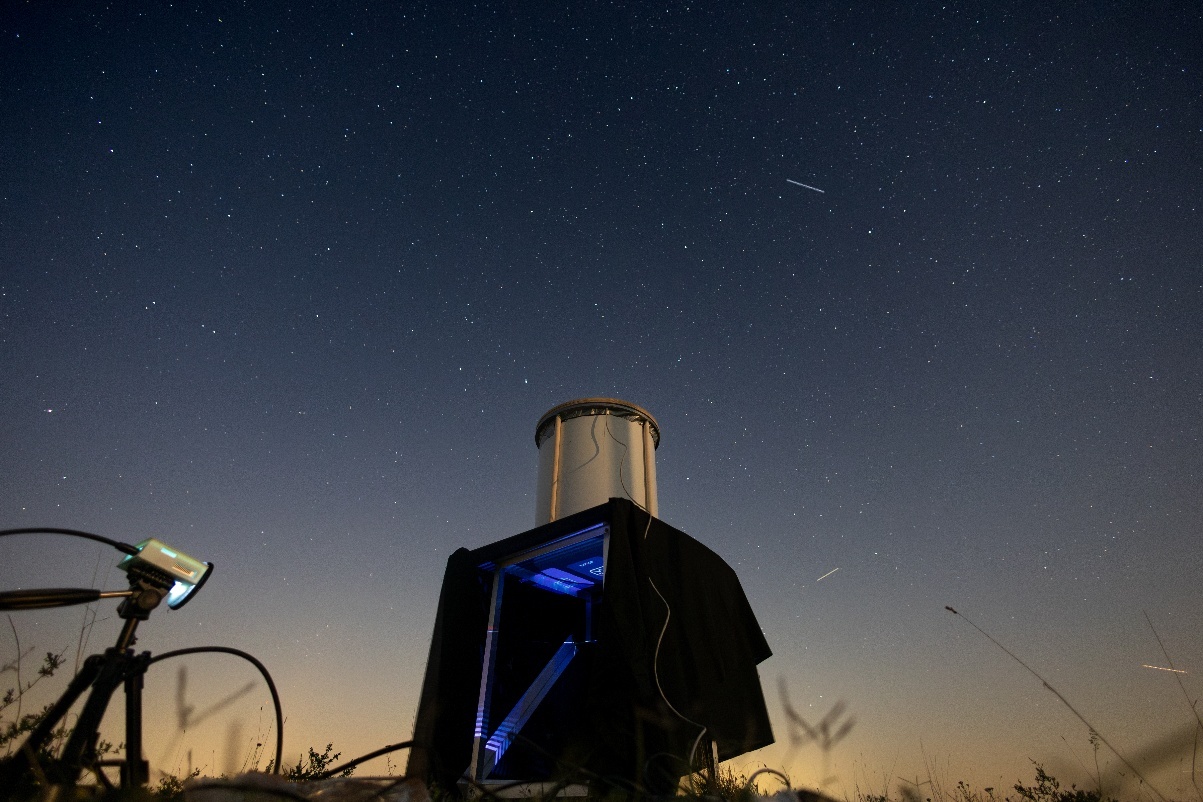 |
| **D** |  |  |
| 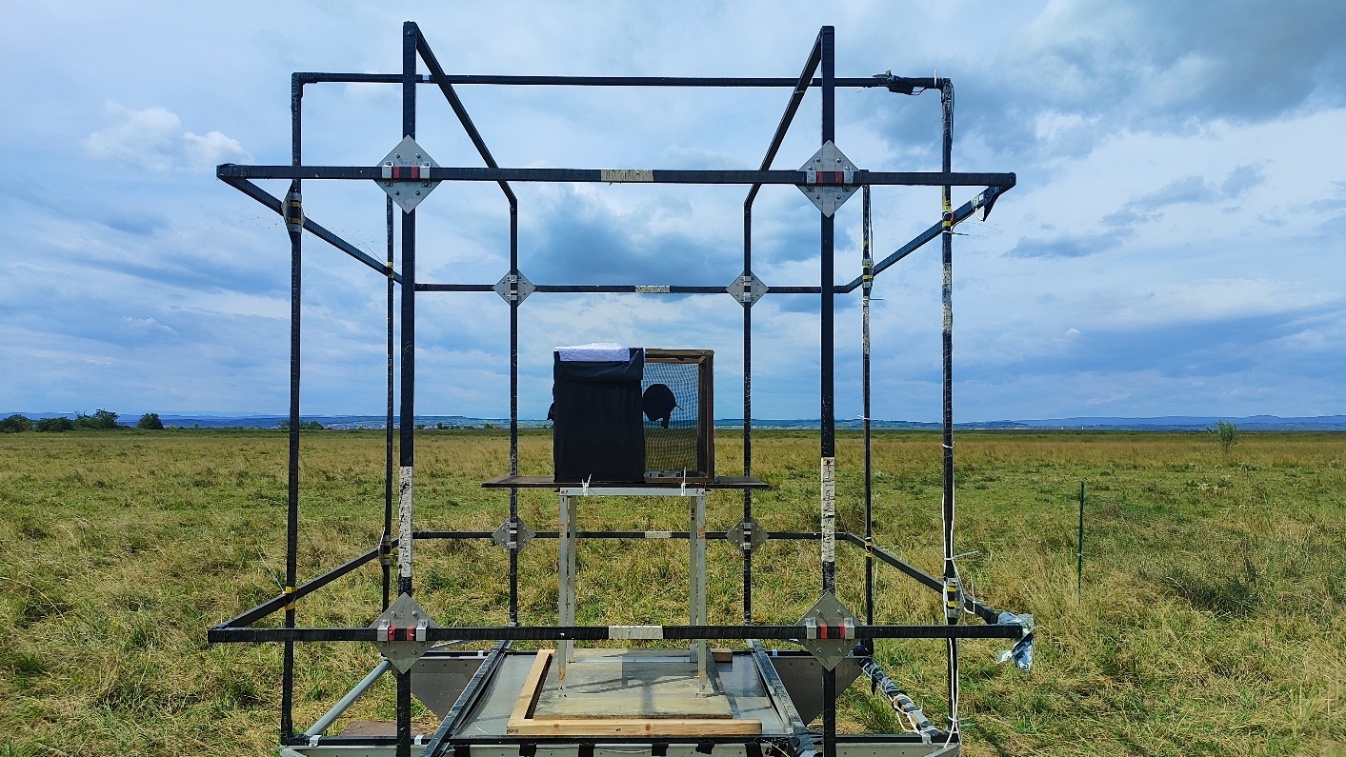 |  | 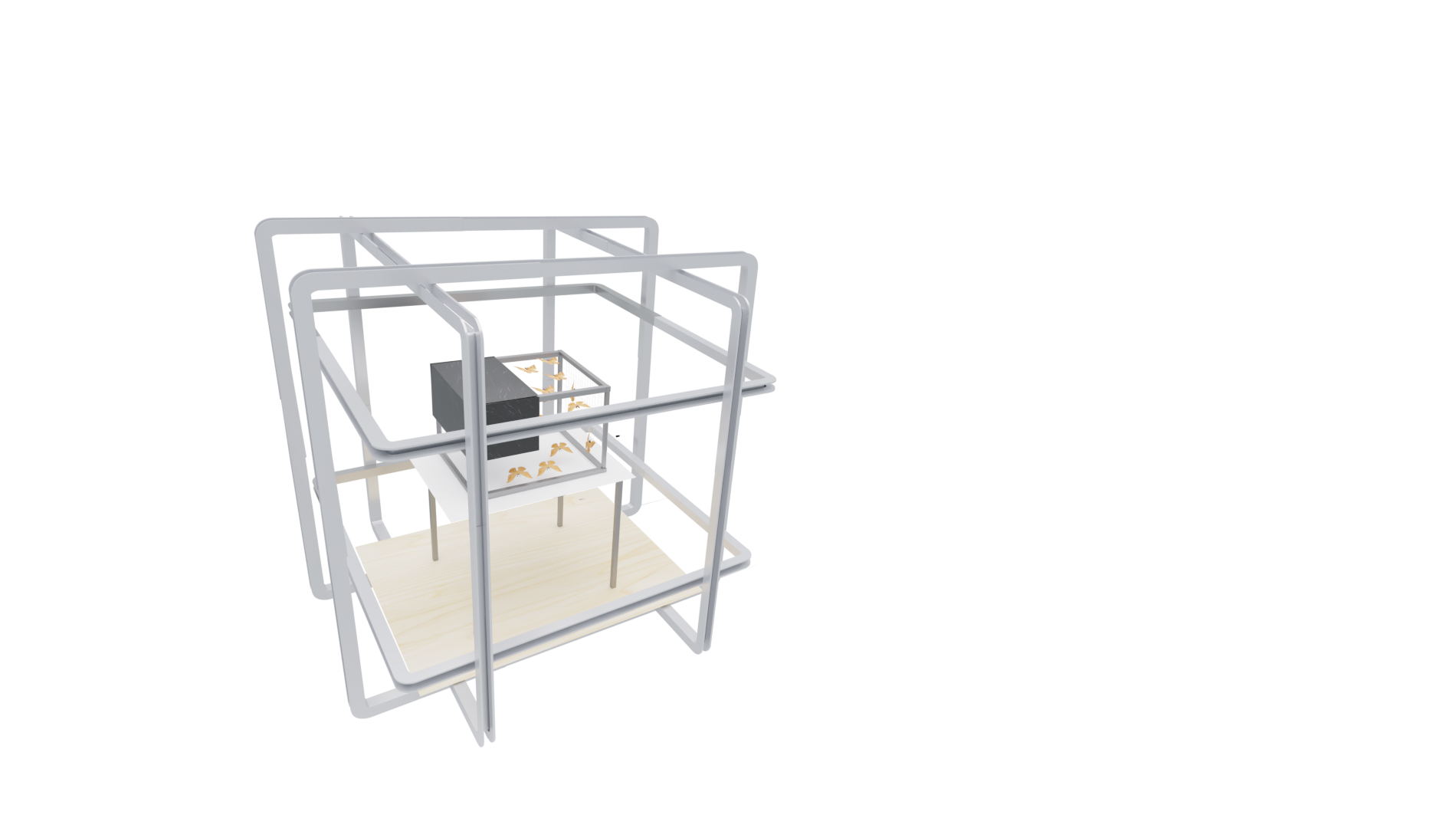 |

**Figure S2. Activity pattens of different red underwings.**

| 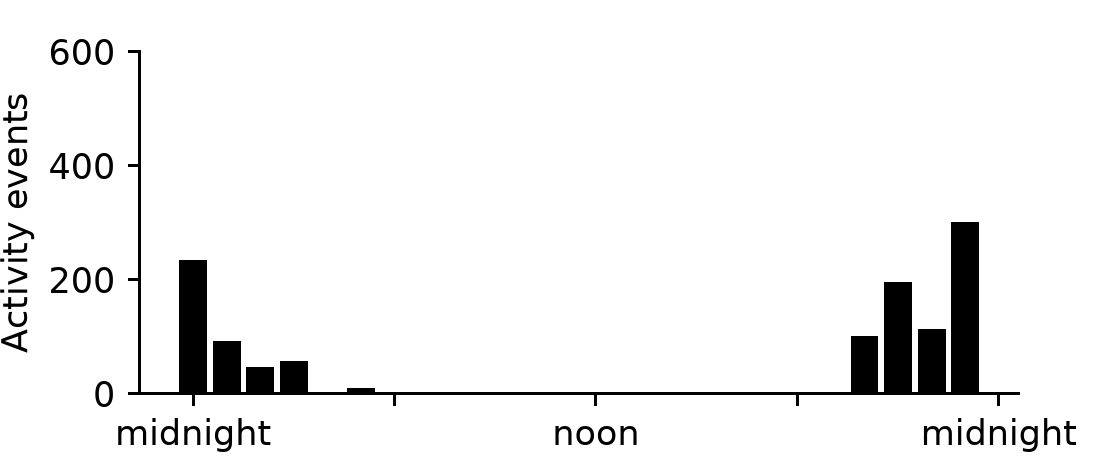 | 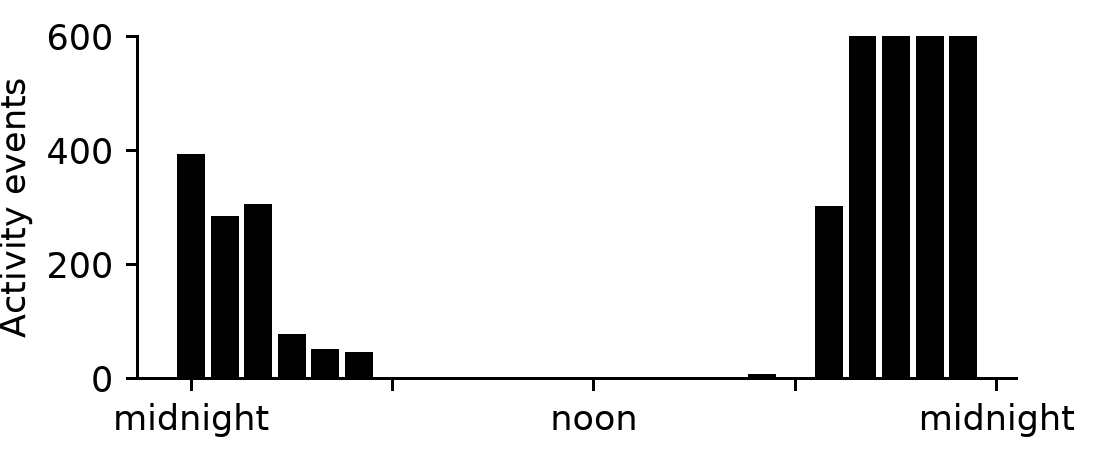 |
| --- | --- |
| 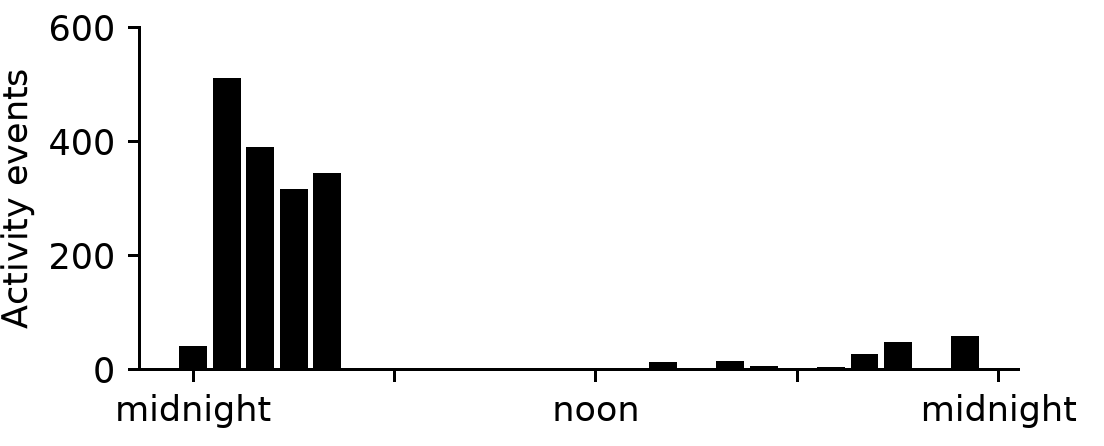 | 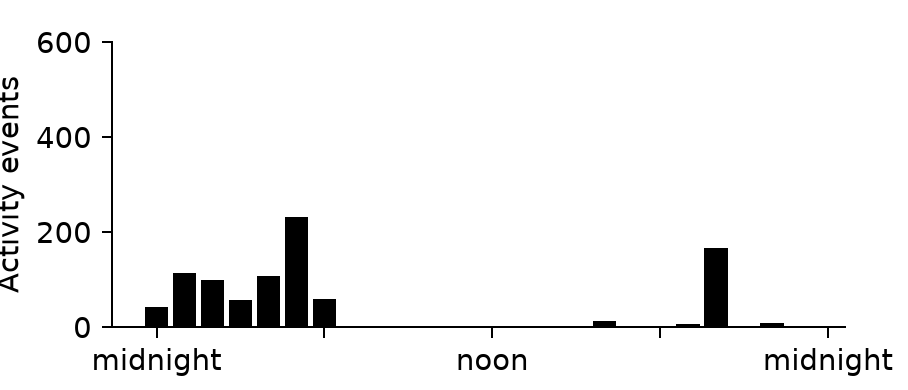 |
| 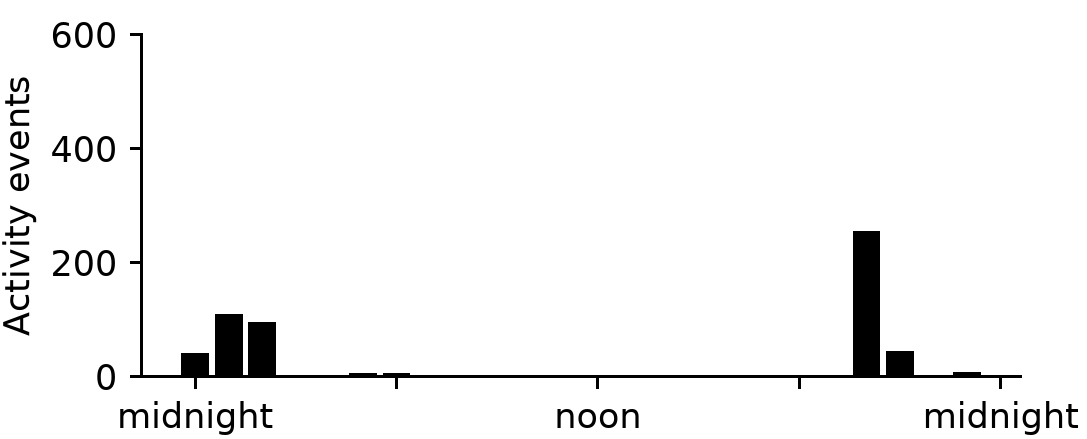 | 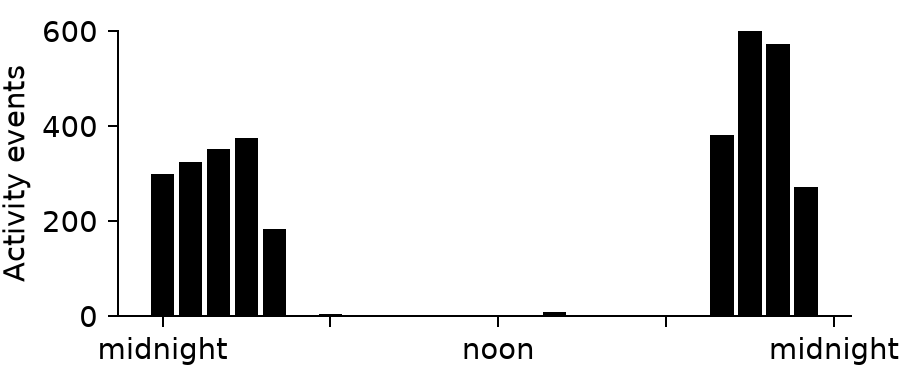 |
| 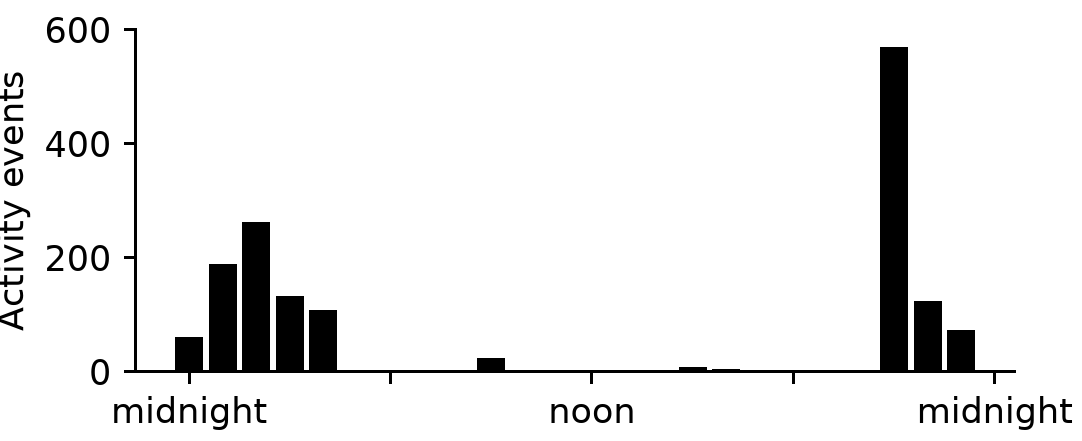 | 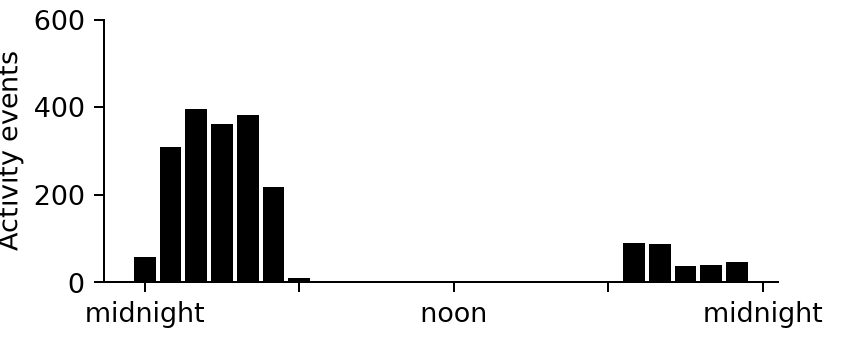 |

**Figures S3. Distribution of red underwings according to data from citizen science projects**

**A** iNaturalist (https://www.inaturalist.org/taxa/116385-Catocala-nupta#map-tab)
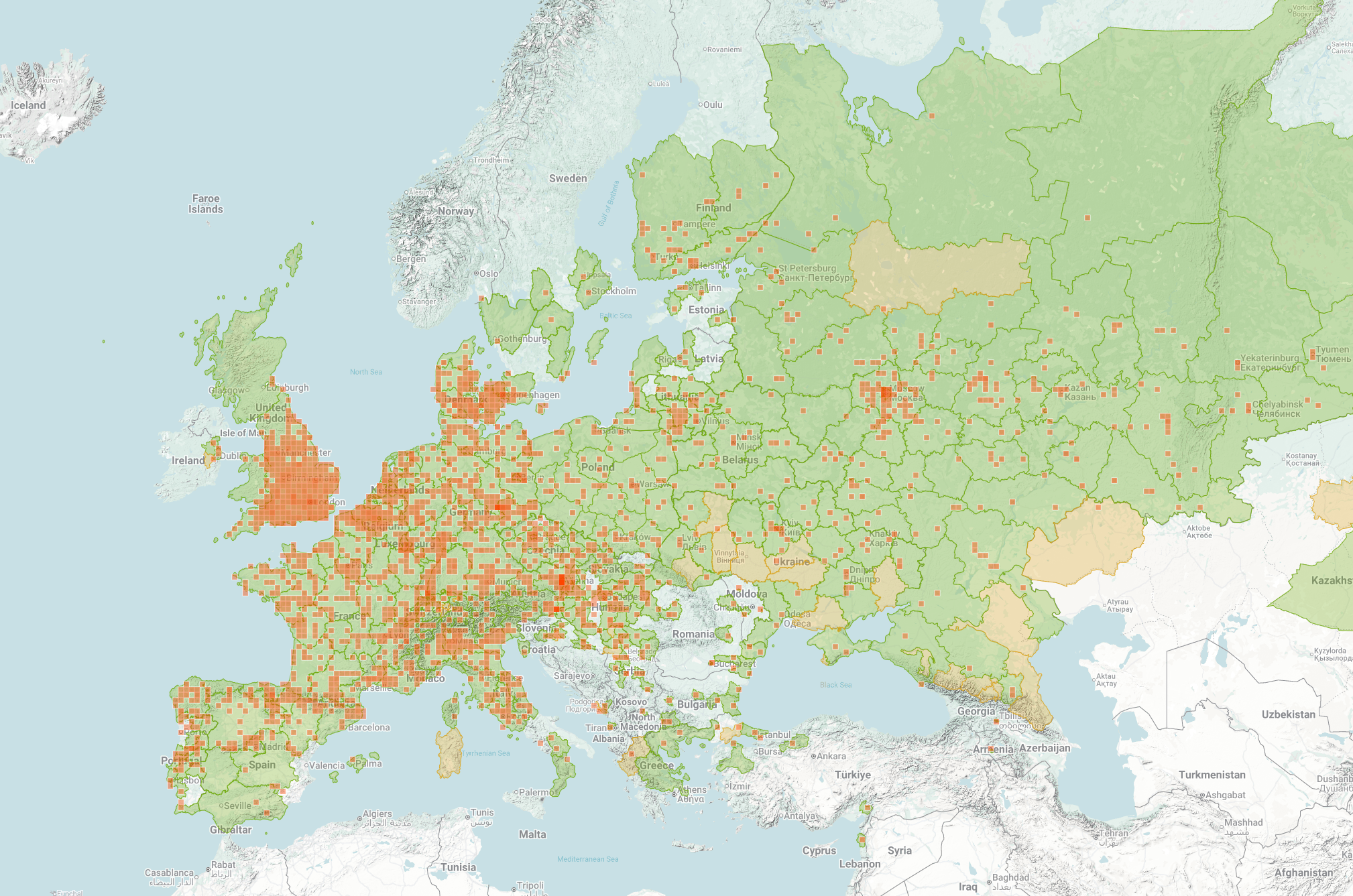

**B** GBIF (https://www.gbif.org/species/1797258)
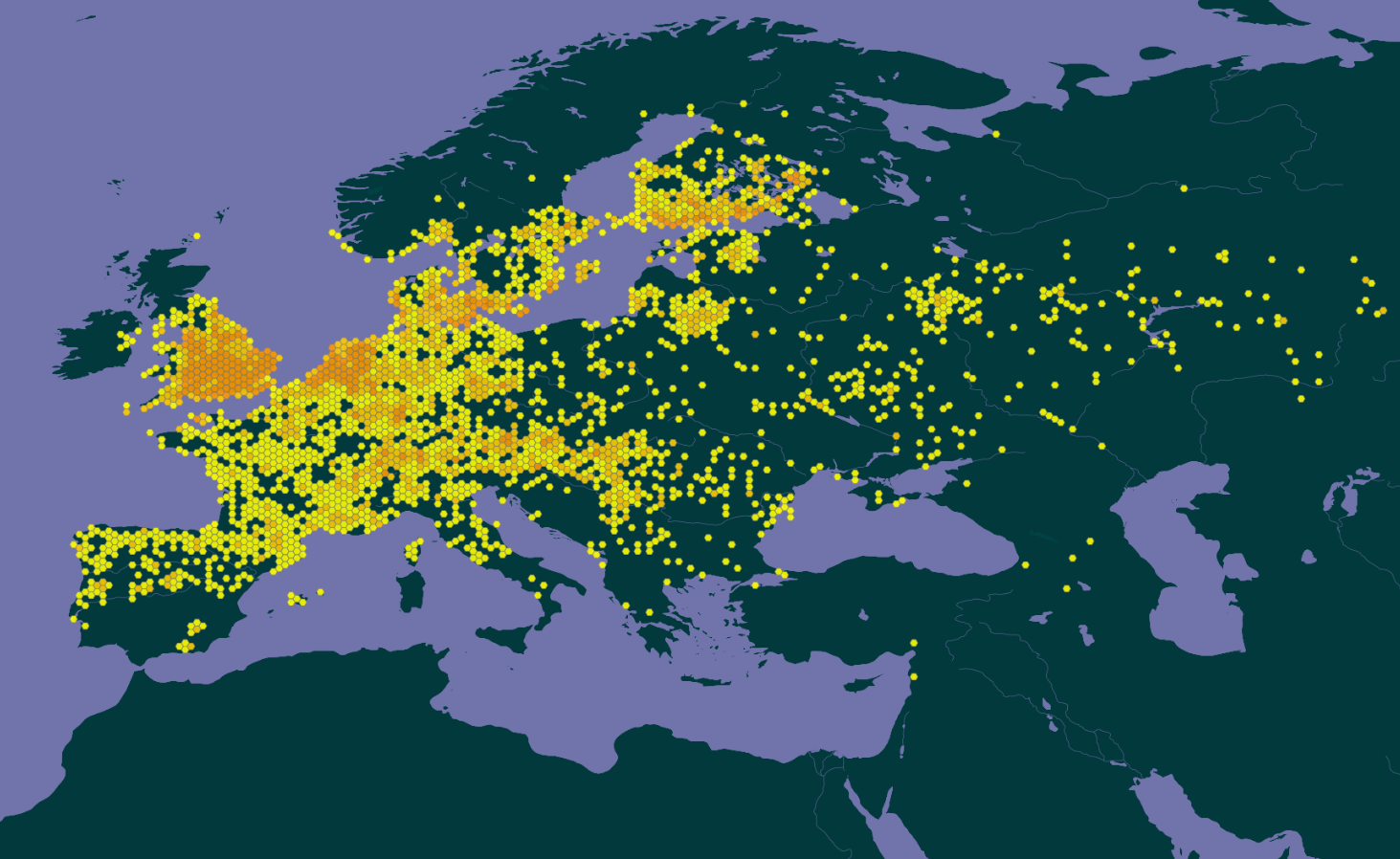

**C** Obsidentify (https://observation.org/species/1792/maps/?start_date=2024-10-06&interval=86400&end_date=2025-10-01&map_type=grid100k)
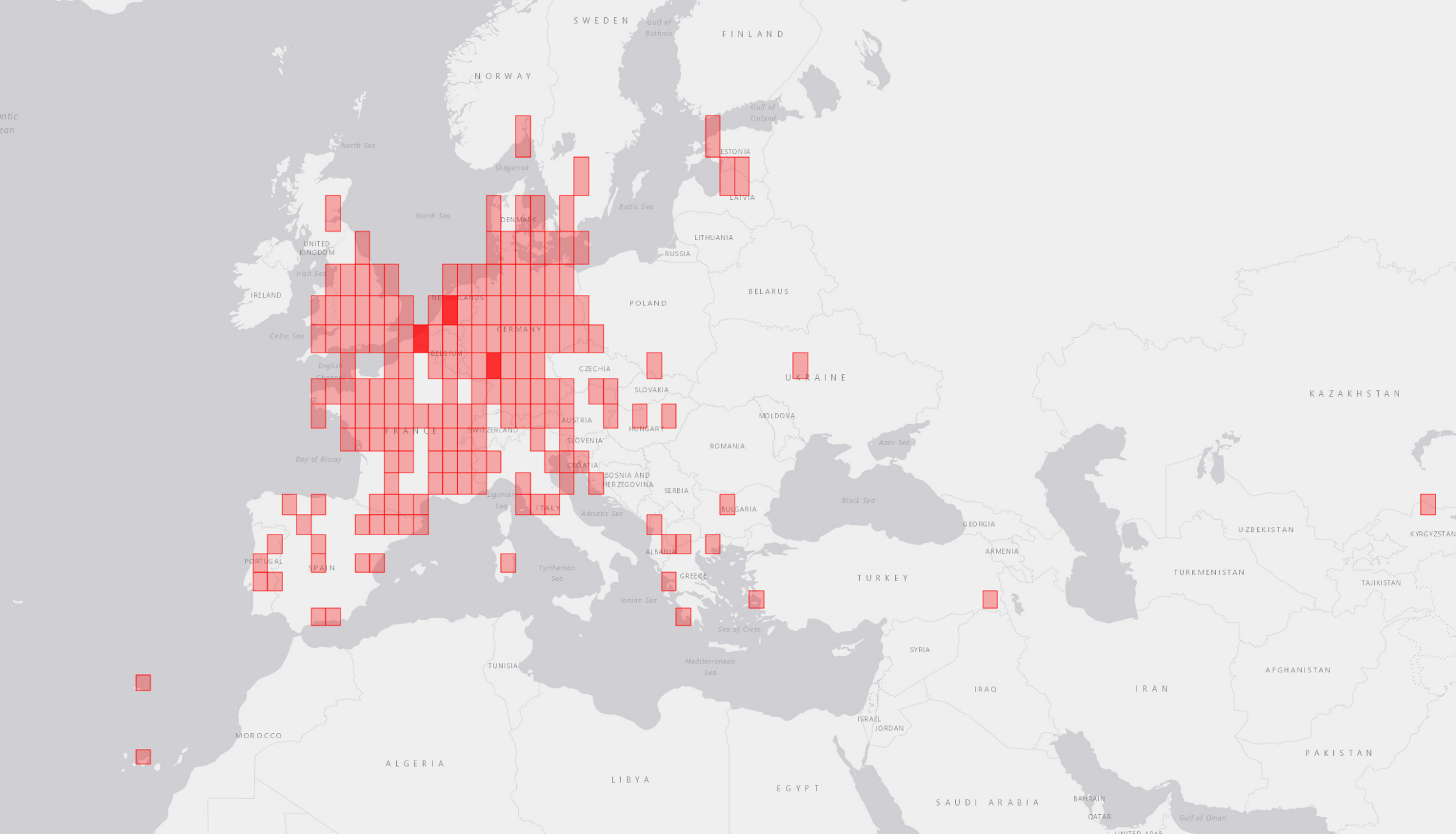

**Figure S4. Possible locations for the virtual magnetic displacement with magnetic parameters used in this study (according to ViMDAL app)**

**A**

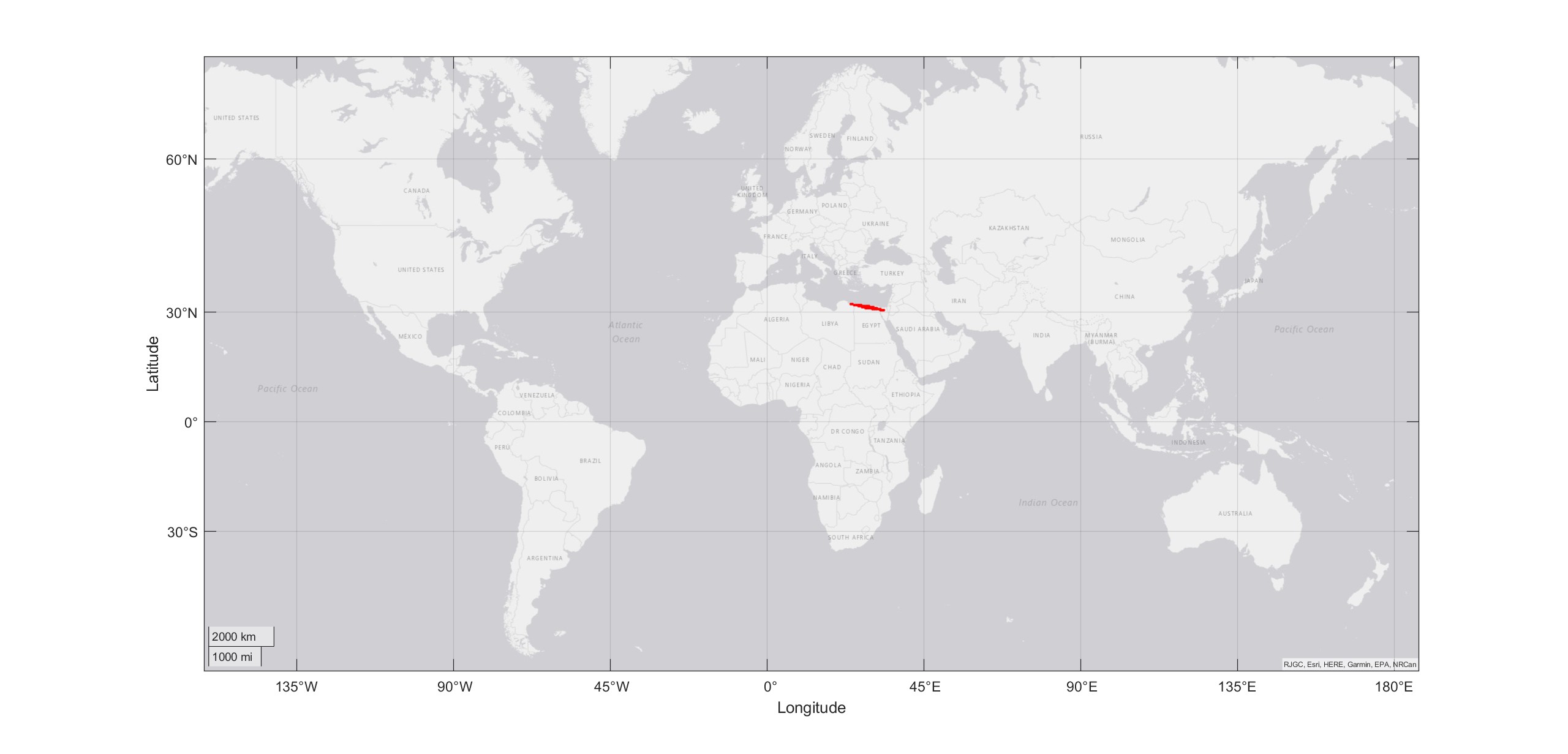

**B**

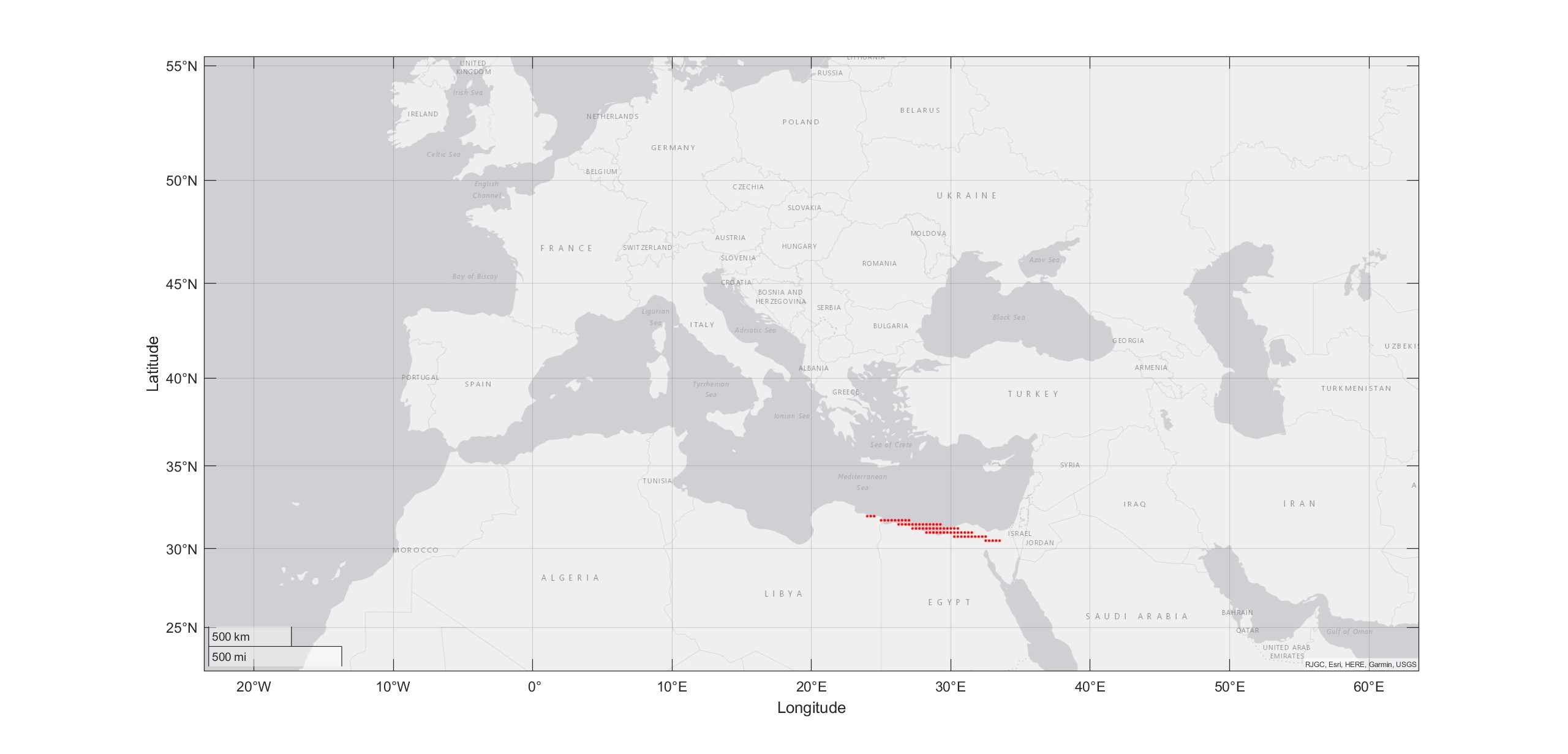

**Table S1. Data of all orientation tests.**

| **Experimental condition** | **№** | **Magnetic cues** | **Celestial cues** | **Mean direction, α** | **Length of mean vector, r** |
| --- | --- | --- | --- | --- | --- |
| **Control condition** | 1 | Natural magnetic field of Illmitz, Austria | Natural starry sky of Illmitz, Austria | 123 | 0.38 |
|  | 2 |  |  | 25 | 0.2 |
|  | 3 |  |  | 209 | 0.6 |
|  | 4 |  |  | 196 | 0.36 |
|  | 5 |  |  | 179 | 0.22 |
|  | 6 |  |  | 253 | 0.33 |
|  | 7 |  |  | 232 | 0.74 |
|  | 8 |  |  | 245 | 0.5 |
|  | 9 |  |  | 178 | 0.2 |
|  | 10 |  |  | 185 | 0.2 |
|  | 11 |  |  | 134 | 0.37 |
|  | 12 |  |  | 239 | 0.2 |
|  | 13 |  |  | 208 | 0.2 |
|  | 14 |  |  | 311 | 0.73 |
|  | 15 |  |  | 273 | 0.24 |
|  | 16 |  |  | 292 | 0.37 |
|  | 17 |  |  | 237 | 0.36 |
|  | 18 |  |  | 189 | 0.35 |
|  | 19 |  |  | 204 | 0.48 |
| **Virtual magnetic displacement** | 1 | Simulated magnetic field of Alexandria, Egypt | Natural starry sky of Illmitz, Austria | 90 | 0.3 |
|  | 2 |  |  | 261 | 0.42 |
|  | 3 |  |  | 135 | 0.33 |
|  | 4 |  |  | 171 | 0.27 |
|  | 5 |  |  | 56 | 0.28 |
|  | 6 |  |  | 83 | 0.36 |
|  | 7 |  |  | 110 | 0.74 |
|  | 8 |  |  | 101 | 0.2 |
|  | 9 |  |  | 137 | 0.36 |
|  | 10 |  |  | 106 | 0.2 |
|  | 11 |  |  | 130 | 0.21 |
|  | 12 |  |  | 189 | 0.77 |
|  | 13 |  |  | 70 | 0.52 |
|  | 14 |  |  | 138 | 0.66 |
|  | 15 |  |  | 215 | 0.59 |
|  | 16 |  |  | 254 | 0.21 |
|  | 17 |  |  | 183 | 0.48 |
|  | 18 |  |  | 251 | 0.4 |
|  | 19 |  |  | 274 | 0.24 |
| **Simulated overcast** | 1 | Natural magnetic field of Illmitz, Austria | No celestial cues (diffusor) | 116 | 0.23 |
|  | 2 |  |  | 233 | 0.25 |
|  | 3 |  |  | 192 | 0.2 |
|  | 4 |  |  | 111 | 0.33 |
|  | 5 |  |  | 259 | 0.26 |
|  | 6 |  |  | 216 | 0.51 |
|  | 7 |  |  | 128 | 0.23 |
|  | 8 |  |  | 94 | 0.35 |
|  | 9 |  |  | 202 | 0.21 |
|  | 10 |  |  | 130 | 0.4 |
|  | 11 |  |  | 304 | 0.21 |
|  | 12 |  |  | 259 | 0.31 |
|  | 13 |  |  | 359 | 0.27 |
|  | 14 |  |  | 305 | 0.96 |
|  | 15 |  |  | 322 | 0.8 |
|  | 16 |  |  | 158 | 0.2 |
| **Vertical magnetic field** | 1 | Vertical magnetic field | Natural starry sky of Illmitz, Austria | 239 | 0.245 |
|  | 2 |  |  | 177 | 0.78 |
|  | 3 |  |  | 202 | 0.8 |
|  | 4 |  |  | 287 | 0.62 |
|  | 5 |  |  | 127 | 0.23 |
|  | 6 |  |  | 144 | 0.21 |
|  | 7 |  |  | 245 | 0.39 |
|  | 8 |  |  | 183 | 0.27 |
|  | 9 |  |  | 249 | 0.3 |
|  | 10 |  |  | 209 | 0.25 |
|  | 11 |  |  | 199 | 0.96 |
|  | 12 |  |  | 159 | 0.2 |
|  | 13 |  |  | 204 | 0.47 |

**Table S2. The results of maximum likelihood estimation for all experimental groups.**

The best fitted models are listed in corresponding column (model names are from Fitak & Johnsen, 2017)

*** -** Best model to describe distribution, **bold –** other probable models to describe distribution (deltaAIC < 2)

**A) Control (before displacement)**

| Model | Description | deltaAIC | AIC_weights |
| --- | --- | --- | --- |
| **M2A*** | **Unimodal** | **0** | **0.53061** |
| **M2B** | **Symmetric modified unimodal** | **1.932** | **0.20199** |
| M2C | Modified unimodal | 2.218 | 0.17503 |
| M5A | Homogenous bimodal | 5.514 | 0.03369 |
| M3B | Symmetric bimodal | 6.347 | 0.02221 |
| M4A | Homogenous axial bimodal | 6.819 | 0.01754 |
| M5B | Bimodal | 8.14 | 0.00906 |
| M4B | Axial bimodal | 8.502 | 0.00756 |
| M1 | Uniform | 11.643 | 0.00157 |
| M3A | Homogenous symmetric bimodal | 13.193 | 0.00072 |
| Best model plots | | | |
| 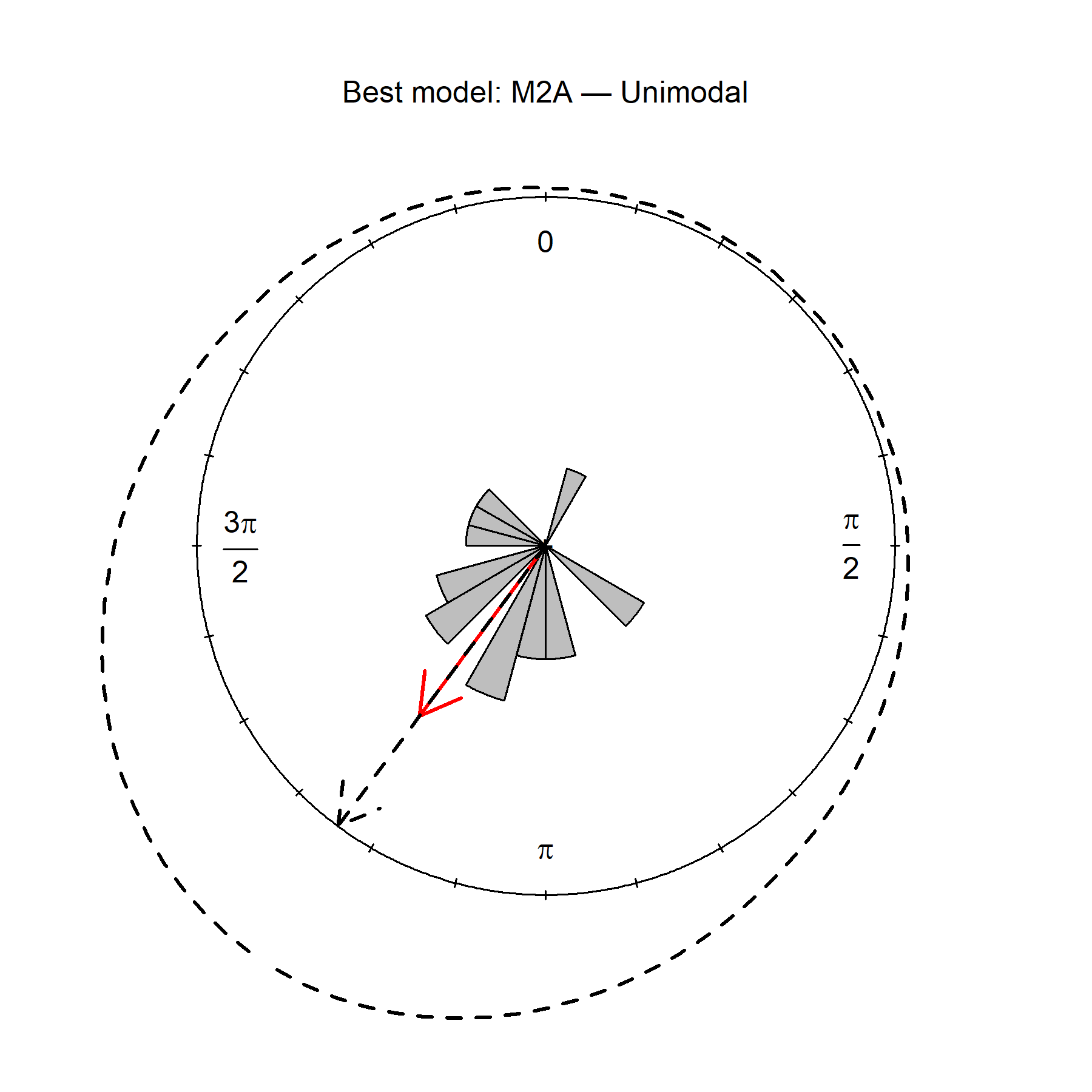 | | | |

**B) After displacement**

| Model | Description | deltaAIC | AIC_weights |  | |
| --- | --- | --- | --- | --- | --- |
| **M5A*** | **Homogenous bimodal** | **0** | **0.41169** |  |  |
| **M5B** | **Bimodal** | **1.582** | **0.18662** |  |  |
| **M2A** | **Unimodal** | **2.088** | **0.14496** |  |  |
| M2B | Symmetric modified unimodal | 2.859 | 0.09855 |  |  |
| M2C | Modified unimodal | 4.632 | 0.04061 |  |  |
| M3B | Symmetric bimodal | 4.867 | 0.03612 |  |  |
| M4A | Homogenous axial bimodal | 4.873 | 0.036 |  |  |
| M1 | Uniform | 6.423 | 0.01659 |  |  |
| M3A | Homogenous symmetric bimodal | 6.589 | 0.01527 |  |  |
| M4B | Axial bimodal | 6.822 | 0.01359 |  |  |
| Best model plots | | First group (> 7 days between control and displacement) | | | Second group (<5 days between control and displacement) |
| 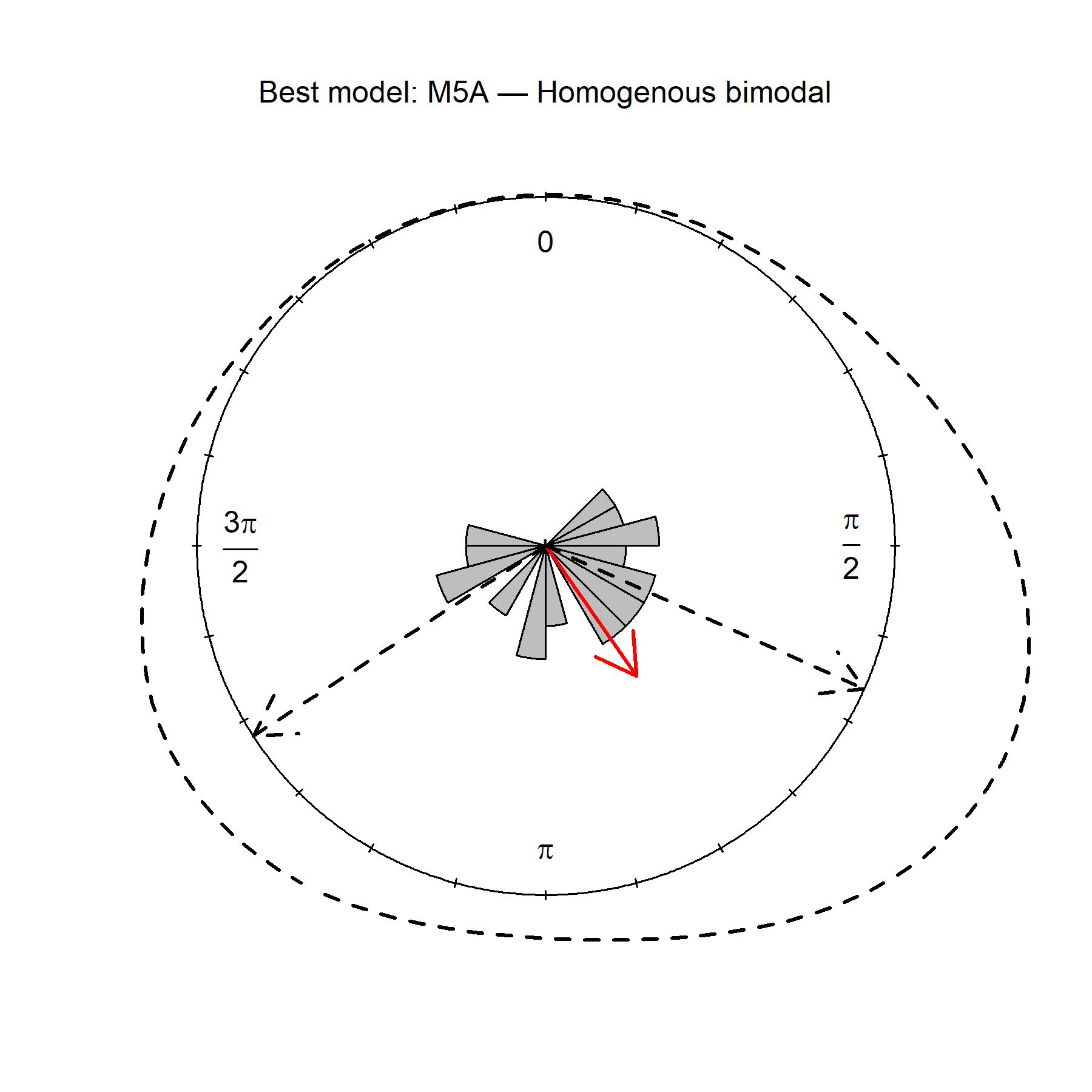 | | 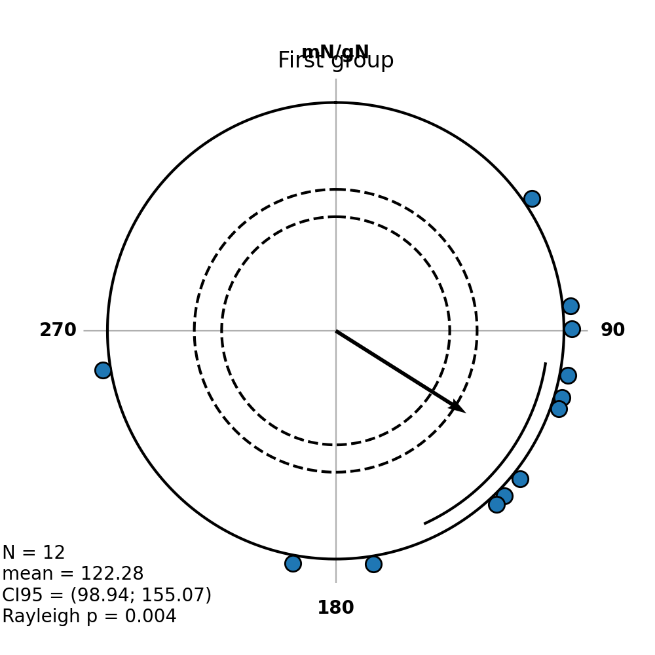 | | | 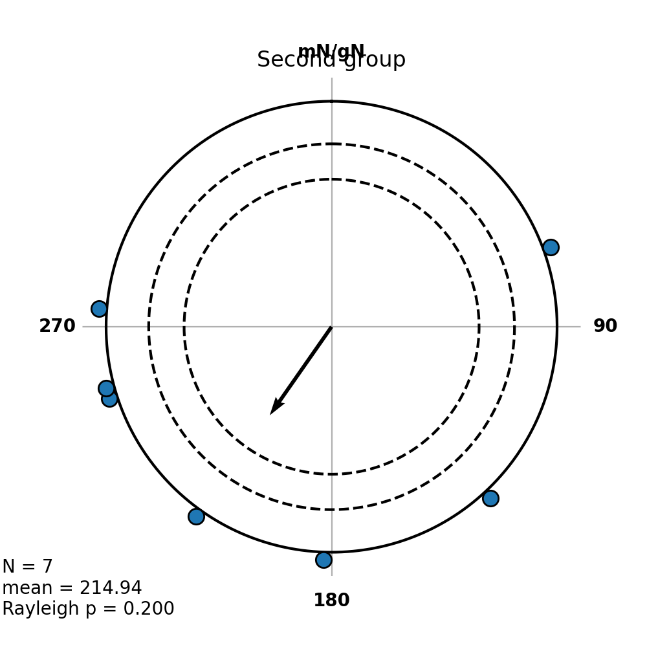 |

**C) Overcast**

| Model | Description | deltaAIC | AIC_weights |
| --- | --- | --- | --- |
| **M1*** | **Uniform** | **0** | **0.32848** |
| **M2A** | **Unimodal** | **1.867** | **0.12916** |
| M2B | Symmetric modified unimodal | 2.345 | 0.10172 |
| M5B | Bimodal | 2.345 | 0.10171 |
| M3A | Homogenous symmetric bimodal | 2.398 | 0.09901 |
| M4B | Axial bimodal | 3.347 | 0.06162 |
| M5A | Homogenous bimodal | 3.689 | 0.05195 |
| M3B | Symmetric bimodal | 3.929 | 0.04607 |
| M2C | Modified unimodal | 4.078 | 0.04275 |
| M4A | Homogenous axial bimodal | 4.338 | 0.03753 |
| Best model plots | | | |
| 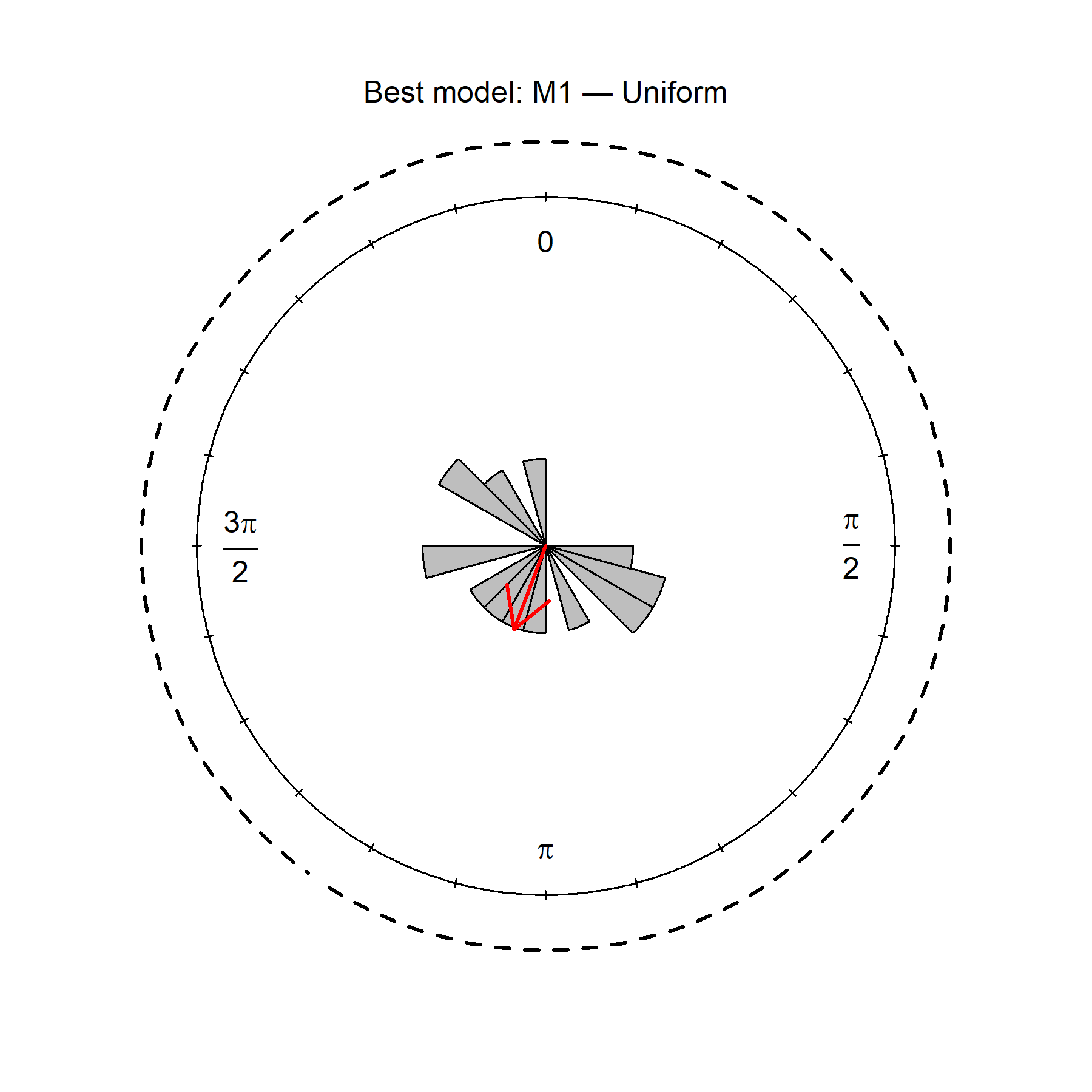 | | | |

**D) VMF**

| Model | Description | deltaAIC | AIC_weights |
| --- | --- | --- | --- |
| **M2A*** | **Unimodal** | **0** | **0.75938** |
| M5A | Homogenous bimodal | 3.264 | 0.14851 |
| M2B | Symmetric modified unimodal | 5.491 | 0.04875 |
| M3B | Symmetric bimodal | 7.831 | 0.01514 |
| M2C | Modified unimodal | 7.99 | 0.01398 |
| M4B | Axial bimodal | 9.324 | 0.00717 |
| M4A | Homogenous axial bimodal | 10.476 | 0.00403 |
| M5B | Bimodal | 12.113 | 0.00178 |
| M1 | Uniform | 13.404 | 0.00093 |
| M3A | Homogenous symmetric bimodal | 15.531 | 0.00032 |
| Best model plots | | | |
| 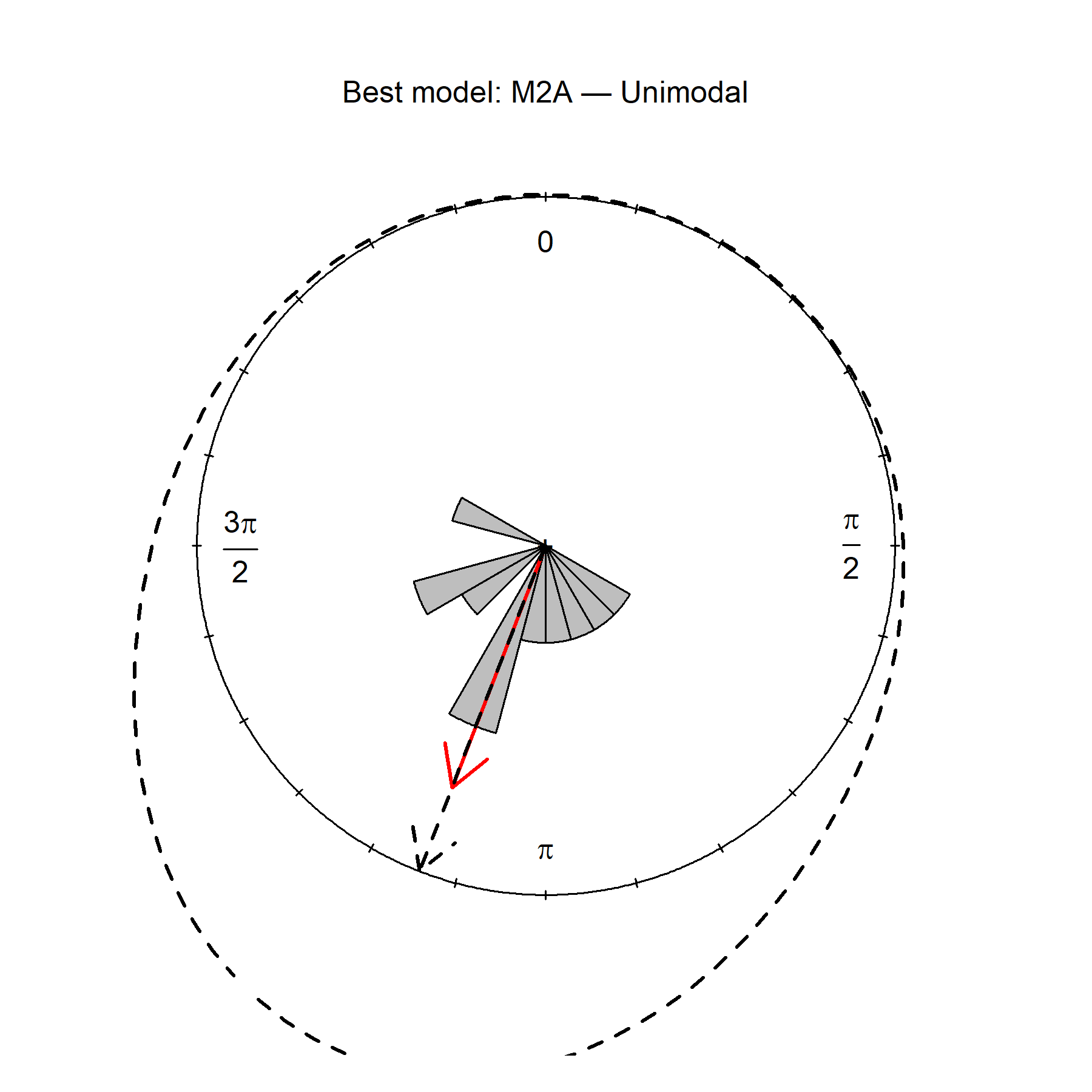 | | | |

**Figure S5. Results of bootstrap analysis.**

Each diagram represents a distribution of lengths of mean vectors that were calculated using a bootstrap technique (n = 100000, see details in the main text of manuscripts, Materials and methods section): A) red underwings tested under overcast conditions vs control group; B) red underwings tested under overcast conditions vs moths in VMF. Vertical blue and red lines indicate 95 and 98 % quantiles, respectively. The red curve is a normal distribution, an orange dot is a length of the mean vector of group in each experimental condition

| **A** | **Control vs Overcast**  > 95% quantile    0.34 - 0.79  > 99% quantile  0.27 - 0.85 | 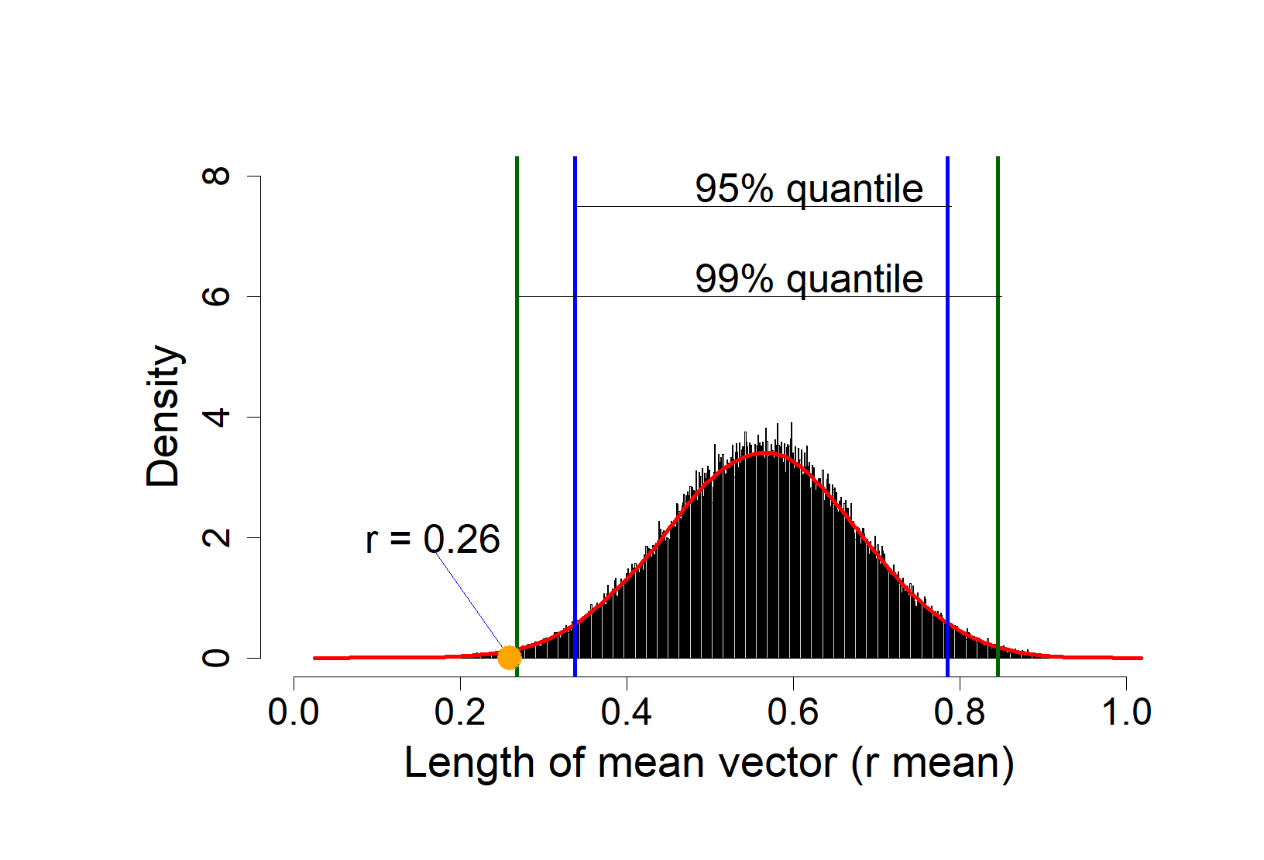 |
| --- | --- | --- |
| **B** | **VMF vs Overcast**  > 95% quantile    0.61 - 0.88  > 99% quantile  0.56 - 0.91 | 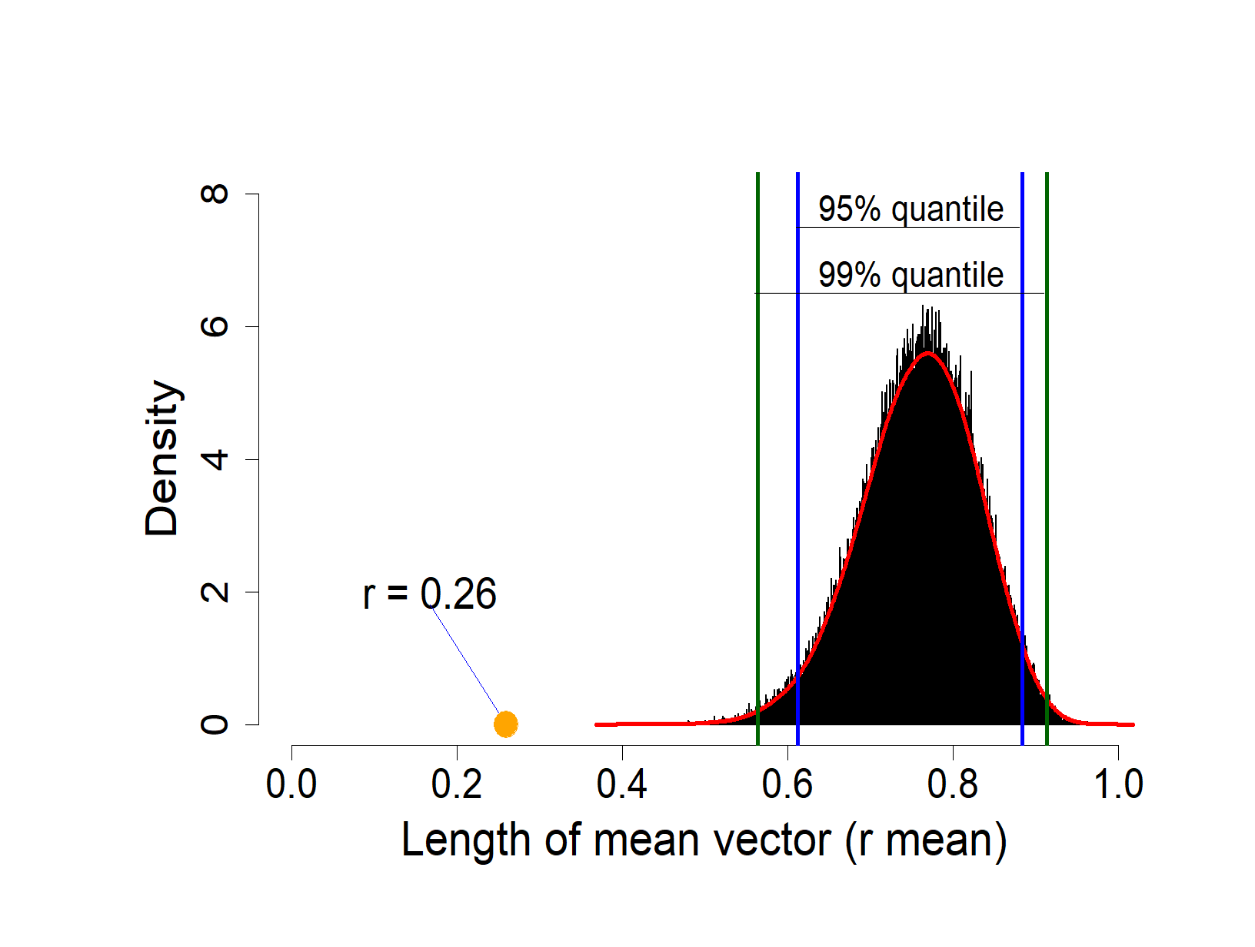 |

**Figure S6. The orientation angles of moths tested with access to clear starry sky plotted as a function of the experimental time.**

Linear regressions of the particular datasets of angles are represented by the orange (moth tested in NMF and under clear sky), black (all moth) and red (moth tested in VMF and under clear sky) lines.

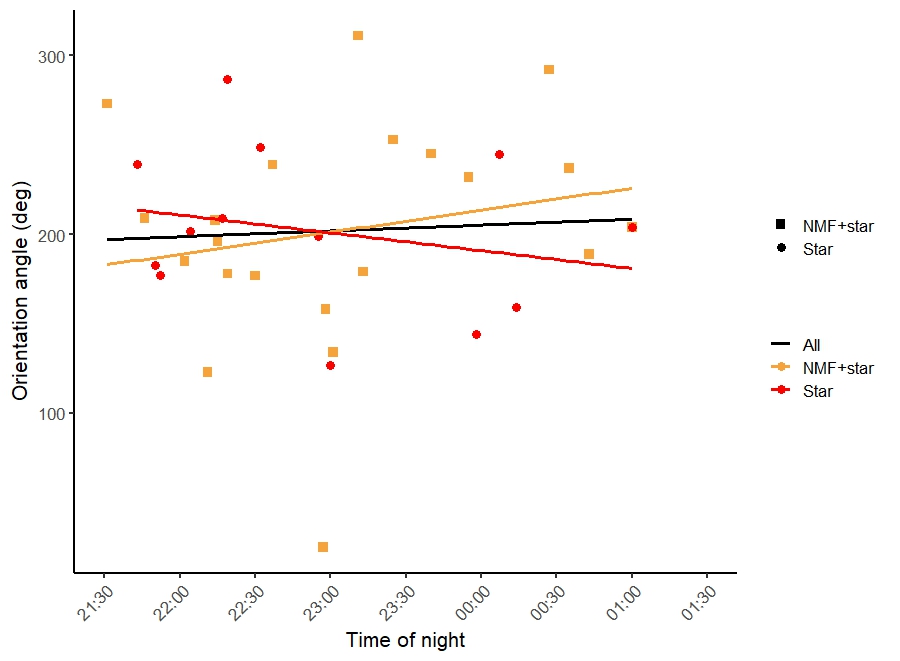
